# Global patterns, drivers and impacts of metabolic traits across amniotes

**DOI:** 10.1101/2024.09.28.615629

**Authors:** Qi Yang, Ziyi Wang, Liqing Fan, Dehua Wang, Fumin Lei, Ying Xiong

## Abstract

Metabolism and climate are thought to influence species richness and diversification, yet few studies have examined whether global patterns of metabolic traits are linked to climate and diversification rate. Here we investigate the drivers of global metabolic rates and their impacts on biodiversity across 2,633 amniote vertebrates. We found consistent distribution patterns of metabolic rates in terrestrial amniotes and marine birds, with metabolic rates increasing with distance from the equator across latitudes. Temperature is a primary climatic variable affecting metabolic rates in birds and mammals, whereas precipitation dominates in terrestrial reptiles. Furthermore, elevated metabolic rates promote diversification rates and reduce extinction risks in birds and terrestrial mammals. Conversely, high metabolic rates decrease diversification rates and increase extinction risks in reptiles. Our results show that global patterns of metabolic rates are driven by various climate variables and may shape diversification and extinction patterns among amniotes in the context of climate change.

## Introduction

Understanding how intrinsic biotic processes influence species diversification and extinction is a fascinating and complex topic in biodiversity conservation (*1*). Metabolism, which provides the energy required for all biological processes—such as reproduction, growth, and generation length—plays a pivotal role in shaping the distribution and diversity of life on Earth (*2-6*). First, thermogenic metabolism indicates thermal tolerance limits for distributional shifts in response to temperature changes. It suggests that several species with broader metabolic tolerances can expand into regions of higher elevation or latitude, whereas those with limited tolerance experience significant population declines as global climate becomes warming (*7-11*). Second, higher metabolic rates are associated with increased species richness at higher latitudes and reduced extinction risks. For example, marine endotherms exhibit peak diversity in polar regions, and island endotherms with elevated metabolic rates face lower extinction risks compared to ectotherms (*4, 12*). Therefore, climate variables are the prevailing explanations for species richness and diversification on a large scale (*13*), but physiological metabolism can also play a significant role in influencing these patterns.

The metabolic rate is a fundamental indicator of metabolic traits, varying according to body mass, temperature, and species thermoregulation (*2*). Temperature presents several challenges to fuel the metabolic demands for physiological performance, particularly concerning metabolic rate and body mass related to regulation of body temperature (*2, 14, 15*). Metabolic rates in ectotherms increase with temperature to improve their environmental fitness, while additional metabolic costs are more vulnerable to impact ecological survival of these species (*16, 17*). Moreover, in accordance with Bergmann’s rule regarding the surface-to-volume ratio of organisms, the body size of endotherms tends to increase in lower temperatures to reduce heat loss at higher latitudes or elevations (*18-20*), but it remains unclear well whether metabolic rate predictably decreases with geographic gradients in temperature. Therefore, exploring the global patterns, drivers, and impacts of metabolic traits across amniotes has profound impacts on understanding the mechanisms underpinning temperature adaptation and biodiversity.

Here we examined global patterns and drivers of metabolic rates across 2,633 amniote species and evaluated the impact of these metabolic rates on species diversification and extinction risk using a hypothesized framework (Fig. 1). We tested the predictable relationship between temperature and metabolic rates through spatial autocorrelation regression (Fig. 1A). By integrating spatial structural equation models, we estimated both the direct and indirect effects of metabolic rates on diversification rates, as well as extinction risks, across terrestrial and marine amniotes (Fig. 1B, C). This study provides robust evidence for a fundamental role of metabolic rates in shaping species diversification and extinction on a global scale.

**Fig. 1.**
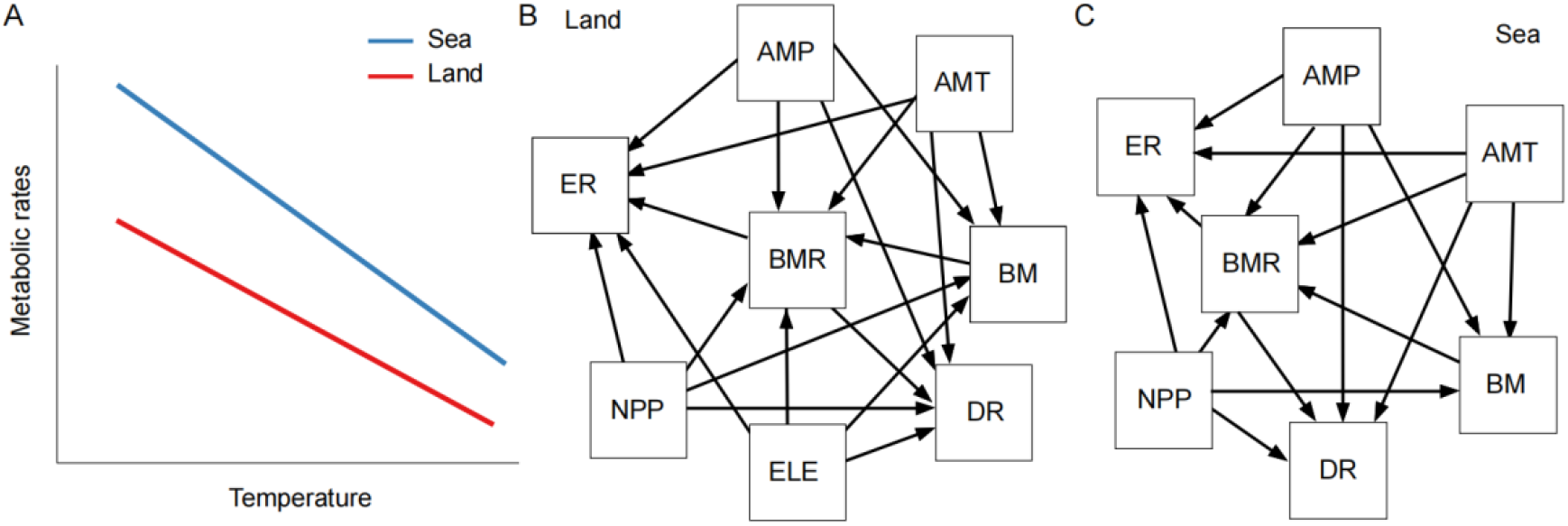
Hypothesized cause and effect among metabolic trait, biodiversity and environmental variables. **A**. Hypothesized relationships between temperature and metabolic rate. Hypothesized structural equation models in terrestrial amniotes (**B**) and marine amniotes (**C**). Abbreviations: AMT, annual mean temperature; AMP, annual mean precipitation; NPP, net primary productivity; ELE, elevation; BMR, basal metabolic rate; BM, body mass; DR, diversification rate; ER, extinction risk.

## Results and Discussion

We found consistent patterns of metabolic rates among terrestrial amniotes (Fig. 2, Figs. S1, S2). Metabolic rates were lowest in tropical regions and exhibited a clear increase along latitudinal gradients toward the poles. These patterns indicate that metabolic rates reflect hemispheric asymmetry, with the northern hemisphere exhibiting higher metabolic rates compared to the southern hemisphere. The American continent was an exception for reptiles, as it displayed higher metabolic rates in the southern regions (Fig. 2C, Figs. S1C, S2C). Moreover, the Qinghai-Tibet Plateau exhibited a distribution of high metabolic rates due to low ambient temperature, despite its low latitude. However, the patterns differed among marine amniotes. Marine birds exhibited a similar pattern to that of terrestrial birds, while no clear pattern was observed in marine mammals and reptiles (Fig. 2, Figs. S1, S2). Some studies have reported that global patterns of body size in birds and marine mammals are related to temperature adaptation (*20-22*), following Bergmann’s rule. However, variations in metabolic rates are likely influenced by additional factors in the marine environment, such as higher thermal conductivity and lower oxygen content compared to terrestrial ones (*23, 24*).

**Fig. 2.**
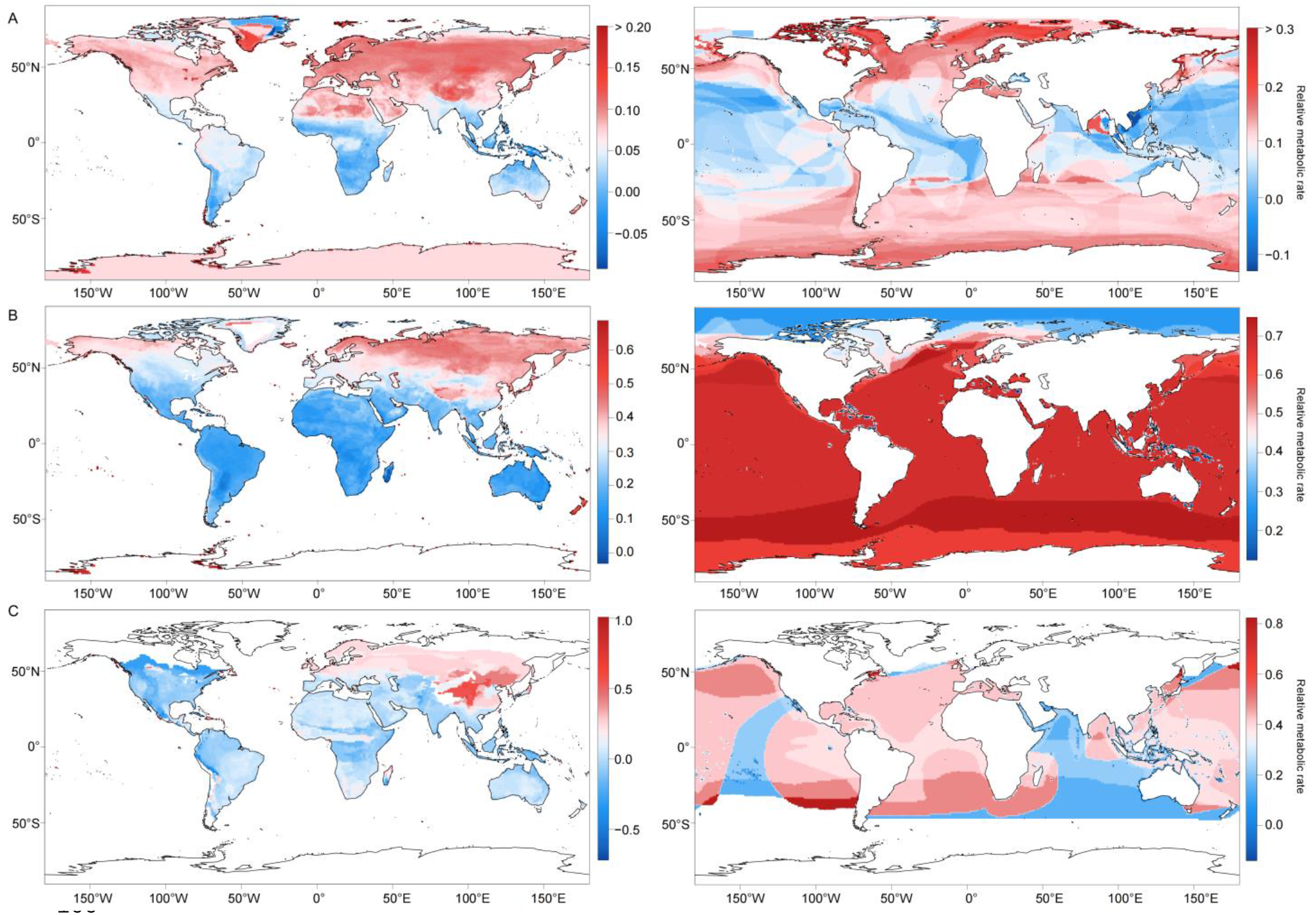
Global patterns of metabolic traits among terrestrial and marine amniotes. Relative metabolic rates increase with distance from the equator across latitudes in terrestrial birds (**A**), mammals (**B**), and reptiles (**C**), while a different pattern is found among marine amniotes.

We used spatial autocorrelation regression models (SARMs) to evaluate the relationships between metabolic rates and variables that may explain metabolic patterns (Fig.3, Tables S1, S2). Single-predictor SARMs revealed that annual mean temperatures (AMT) were significantly negatively correlated with metabolic rates in birds, terrestrial mammals and reptiles (Fig. 3A, Figs. S3, S4, Tables S1, S2). This can likely be attributed to the thermoregulatory types, which birds and mammals, as endotherms, increase heat production through elevated metabolic rates in cold ambient temperatures (*25, 26*). However, this relationship in terrestrial reptiles contradicts the principles of ectothermic physiology due to the potential effect of body mass (*16*), as mass ^2/3^-specific metabolic rate is positively related with temperature (Fig. S3C, Tables S1). Moreover, marine mammals demonstrate a positive relationship between temperature and metabolic rates (Fig.3A, Fig. S4B, Table S2), which may be associated with lower metabolic costs for swimming or thermogenesis during the transition from a terrestrial to an aquatic lifestyle (*24, 27*). Multivariate SARMs further supported these findings, but we did not test the associations between ambient temperatures and metabolic rates in reptiles due to the effect of other variables (Fig. S5, Tables S1, S2).

**Fig. 3.**
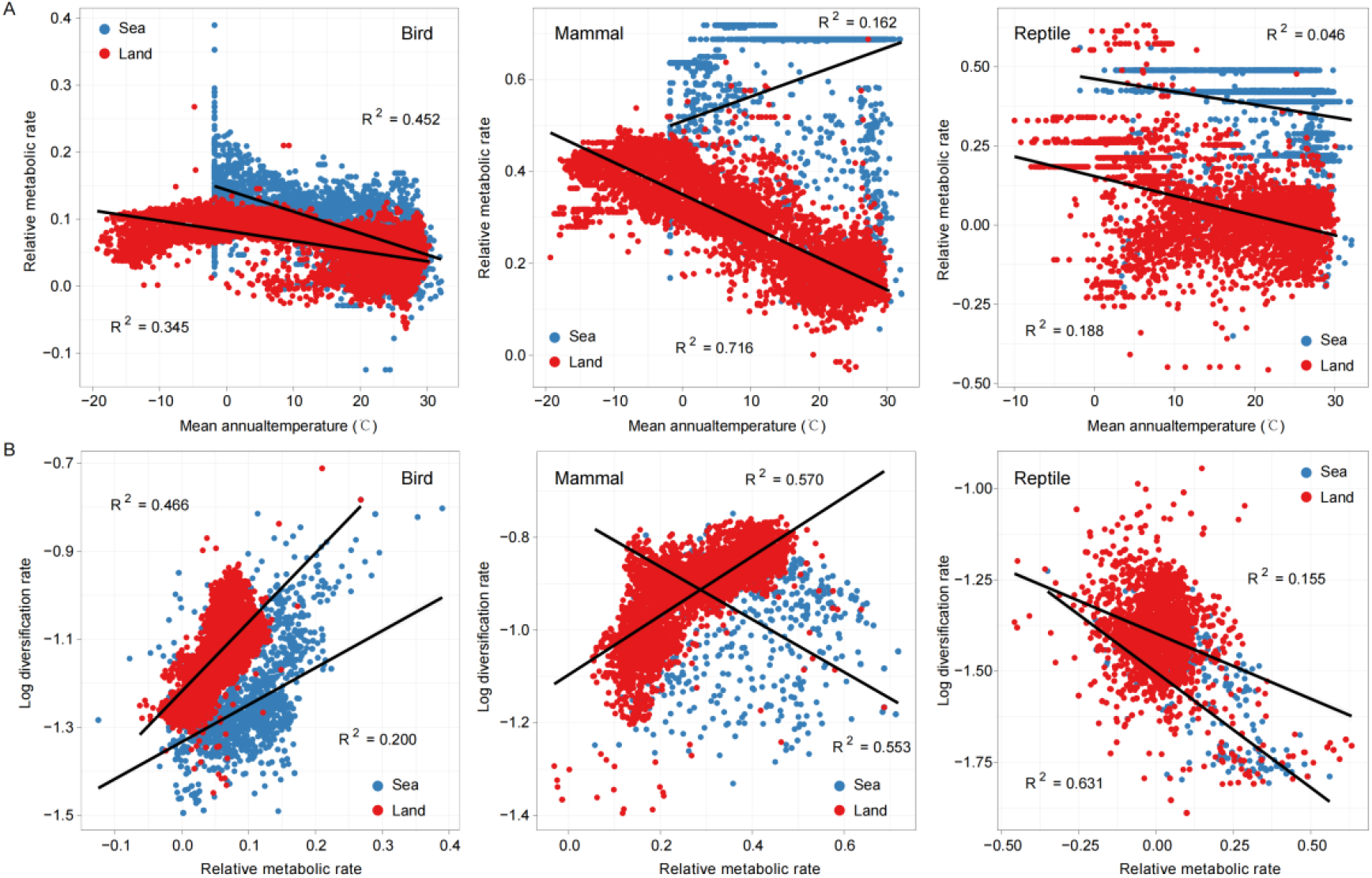
The relationships between temperature and relative metabolic rate, as well as between relative metabolic rate and diversification rate. **A**. Significantly negative relationships between temperature and relative metabolic rate in birds, terrestrial mammals and reptiles, but a positive relationship in marine mammals. **B**. Significantly positive relationships between relative metabolic rate and diversification rate in birds and land mammals, while negative relationships in reptiles and marine mammals.

In addition to ambient temperature, net primary productivity (NPP) showed significantly negative associations with relative metabolic rates across all groups, except for marine endotherms (Tables S1, S2). Although high NPP is believed to enhance biomass accumulation, thereby providing more food for animals (*28, 29*), these relationships suggest that available food resources do not necessarily improve physiological metabolism. By contrast, high NPP is associated with increased metabolic rates in marine mammals (Tables S2), supporting the notion that productivity contributes to body size and species richness in marine mammals (*13, 21*). Furthermore, we found that annual mean precipitation (AMP) is positively correlated with metabolic rates in all terrestrial clades, rather than marine groups (Tables S1, S2). Notably, the metabolic rates of land reptiles demonstrated a stronger correlation with AMP than with AMT (Tables S1). This suggests that ambient aridity may impose greater constraints on metabolism compared to humidity, as an increase in metabolic capacity monotonically increases respiratory water loss (*30*). These findings suggest that ambient temperature plays a consistent and dominant role in shaping metabolic rate patterns in terrestrial birds and mammals, while precipitation dominates in terrestrial reptiles. However, these environmental predictors were not consistently associated with metabolism in marine clades.

Importantly, even after controlling for the effects of body mass, the patterns of metabolic rates were positively correlated with body mass across all groups, with the exception of terrestrial reptiles (Fig. S5, Tables S1, S2). These relationships may be elucidated by the patterns of body mass that correspond to metabolic rates observed (Fig. S6). These relationships were stronger than those associated with environmental variables, which aligns with the metabolic theory of ecology (*2*). These results not only support the traditional heat conservation, but also provide a metabolic mechanism explaining Bergmann’s rule.

Spatial structural equation models revealed a strong positive effect of metabolic rates on diversification rates in birds and terrestrial mammals, while indicating a significantly negative effect in reptiles and marine mammals (Fig. 4, Tables S3, S4). Exogenous environmental predictors exerted both direct and indirect effects on species diversification by influencing metabolic rates. High AMT and NPP increased metabolic rates, thereby promoting diversification in marine mammals (Fig. 4B, Table S4), consistent with the kinetic energy hypothesis (*31, 32*). Inconsistent with the kinetic energy hypothesis, increased metabolic rates induced by low temperatures, rather than high temperatures, enhance the rates of species diversification in birds and terrestrial mammals (Fig. 4A, B, Table S3). Although AMP drove an increased metabolic rate in both terrestrial endotherms and reptiles, it had an inverse effect on diversification rates (Fig. 4B, C, Table S3). This result supports that a high metabolic cost constrains the rates of climatic niche evolution and diversification in reptiles, though all amniotes possess advantages in water regulation (*25*). Unlike marine birds that possess aerial or semi-aquatic lifestyles, AMP had no effect on the metabolic rates of marine mammals and reptiles that have entirely aquatic lifestyles (Fig. 4, Table S4).

**Fig. 4.**
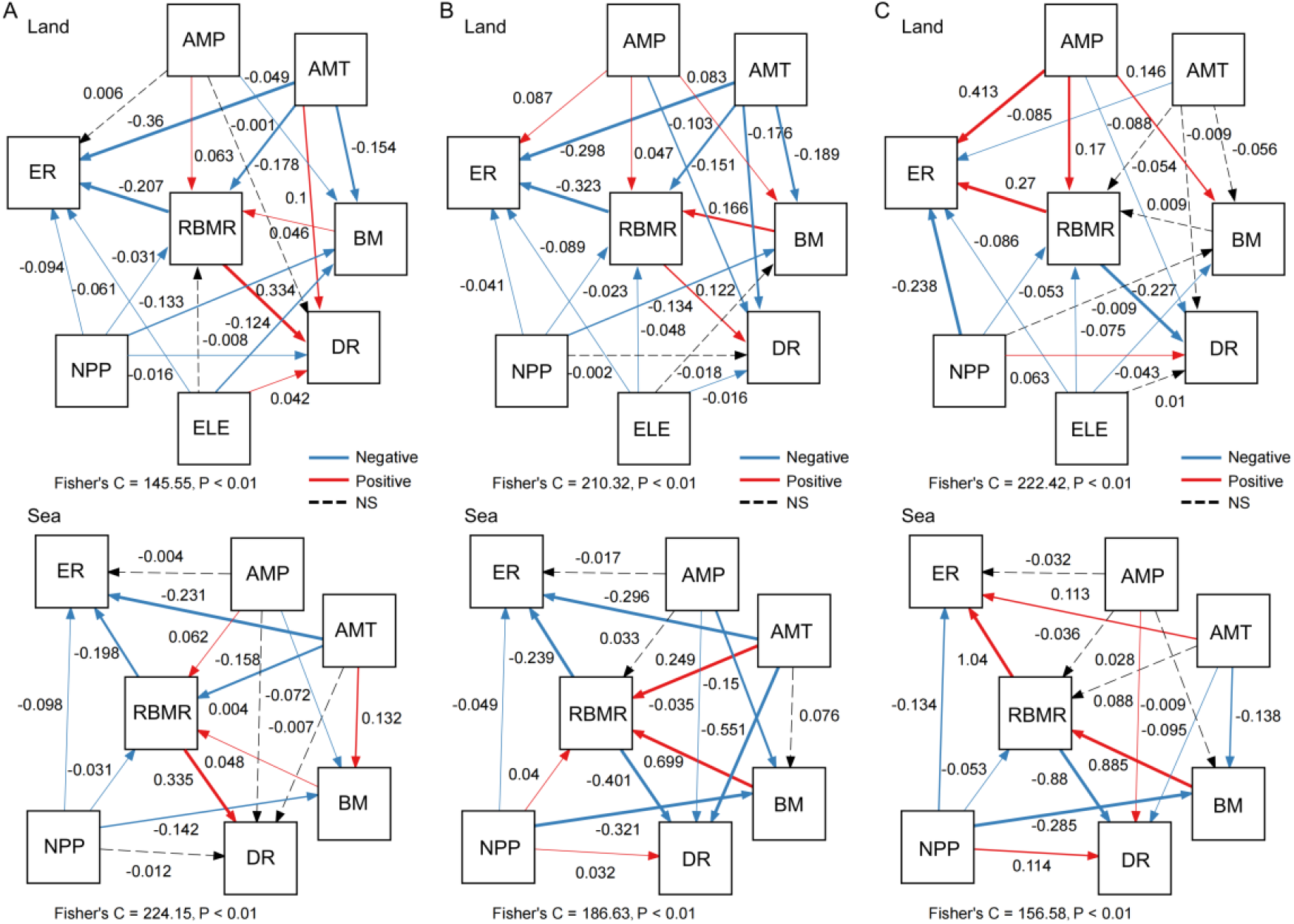
Spatial structural equation models of metabolic rates: drivers and impacts. **A**. In birds. **B**. In mammals. **C**. In reptiles. The values on the arrows represent standardized path coefficients. Red arrows indicate significantly positive effects, while blue arrows indicate significantly negative effects. Non-significant paths are indicated by dashed arrows. The thickness of the path arrows reflects the strength of the relationships. The results of the structural equation models are summarized in Tables S3 and S4. Abbreviations: AMT, annual mean temperature; AMP, annual mean precipitation; NPP, net primary productivity; ELE, elevation; RBMR, relative metabolic rate; BM, body mass; DR, diversification rate; ER, extinction risk.

Similarly, metabolic rates exhibited a significant negative impact on extinction risks in both birds and mammals (Fig. 4A, B, Tables S3, S4). By contrast, elevated metabolic rates were associated with increased extinction risks in reptiles (Fig. 4C, Tables S3, S4). Consistent with our previous report (*4*), metabolic rates have an inverse effect on species extinction in species with different thermoregulatory types. Extinction risks decreased with increasing AMT and NPP across all groups, with the exception of a direct positive effect of AMT on the extinction risks of marine reptiles (Fig. 4, Tables S3, S4). This result is consistent with the ‘productivity-richness’ hypothesis, which posits that high productivity facilitates larger population sizes and consequently reduces the risk of extinction (*33, 34*). However, contrary to previous reports (*35, 36*), high AMT is directly associated with low extinction risks (Fig. 4, Tables S3, S4). This suggests that climate warming may have an indirect effect on species extinction by influencing metabolic rates. Moreover, we noted a direct positive effect of AMP on the extinction risks of terrestrial mammals and reptiles, although this effect was not consistently observed in other groups (Fig. 4, Tables S3, S4).

We also calculated mass-specific metabolic rates using both the 2/3 power and the 3/4 power to scale body mass, but these two exponents have been challenged by numerous studies (*37*). As the scaling exponents for mass-metabolism were found to be 0.661, 0.741, and 0.799 for birds, mammals, and reptiles, respectively, relative metabolic rates are more robust and empirical than these methods (*4*), which likely explains some different results among these three types of metabolic rates (Tables S3, S4).

In conclusion, utilizing an unprecedented dataset of amniote vertebrates that integrates metabolic traits, environmental variables, and diversity data, we identified similar patterns of global metabolic traits in terrestrial amniotes and marine birds. However, no discernible distributional patterns were observed in marine mammals and reptiles. The patterns observed in terrestrial endotherms are primarily associated with temperature, which not only supports the heat conservation hypothesis but also provides a metabolic mechanism to explain Bergmann’s rule (*20-22*). In terrestrial reptiles, precipitation, rather than temperature, predominantly influences metabolic rates, likely due to respiratory water loss (*30*). Furthermore, elevated metabolic rates resulting from low temperatures contribute to diversification rates in birds and terrestrial mammals, which is inconsistent with the kinetic energy hypothesis (*31, 32*). By contrast, reduced metabolic rates appear to facilitate species diversification in reptiles and marine mammals. Although the climatic niche and extinction risks in various thermoregulatory vertebrates have been well studied (*4, 25*), our findings highlight the general principle that high metabolic rates reduce extinction risks in endotherms while increasing them in ectotherms. This study suggests that metabolic traits influenced by climate change may play a significant role in shaping diversification and extinction patterns in the future.

## Methods

### Metabolic data

Basal metabolic rates were compiled from our previous study and other literature (see Data S1) (*4*) and converted to kilojoules per hour (kJ/h) from the oxygen consumption rates using the conversion factor of 1 ml O_2_ = 20.083 J. We adjusted the metabolic rates measured at various temperatures to those at 25°C, following previous studies (*4, 38*). Ultimately, by integrating distributional information, we compiled a comprehensive dataset of metabolic rates from 2,633 species of amniote vertebrates, which includes data on 1,310 birds, 863 mammals, and 460 reptiles.

There is a well-established allometric relationship between metabolic rate and body mass, with the scaling exponent for mass-metabolism ranging from 0.2 to 1.02 (*37, 39*). The 2/3-power law and the 3/4-power law are frequently employed to explain metabolic rates scaled by body mass in ecological studies, including food webs and energy fluxes (*37, 40*). To account for the effects of body mass, we considered three parameters of metabolic rates: metabolic rates scaled by body mass to the 2/3 power (MR0.67), metabolic rates scaled by body mass to the 3/4 power (MR0.75), and the residuals obtained by regressing the log-transformed basal metabolic rates against the log-transformed body masses using phylogenetic generalized least squares (PGLS) analysis. We conducted all downstream analyses using these three scaled metabolic rates. In this study, the scaling exponents were 0.661, 0.741, and 0.799 for birds, mammals, and reptiles, respectively, all of which were inconsistent with both the 2/3 and 3/4 power laws. Therefore, the residuals provided a more robust and empirical basis for scaling body mass (*4*), and we presented the results of the residuals in the main text.

### Distributional data, environmental variables and spatial autoregressive analysis

Geographic distributional data were obtained from the IUCN red list website for reptiles and mammals, and from BirdLife International for birds (download on November 29, 2021) (*41*) (http://www.birdlife.org/). We created maps that represented the following spatial patterns based on 1° × 1° grid cells using use a WGS84 coordinate reference system.

For marine environments, three variables including annual mean temperature of sea surface (AMT), net primary production (NPP) and annual mean precipitation (AMP) were calculated. AMT (°C) was obtained from Geographic Data Sharing Infrastructure of Peking University (http://geodata.pku.edu.cn) (*42*). NPP (units of mg C / m^2^ / day) based on the cafe algorithm was calculated by averaging monthly data from the Ocean Productivity website (http://orca.science.oregonstate.edu/) (*43*). Annual mean precipitation obtained from Global Precipitation Climatology Project (http://research.jisao.washington.edu/data_sets/gpcp) (*44*). For territorial environments, AMT, AMP and elevation of each cell were obtained from layers of climate factors from WorldClim (http://www.worldclim.org) at a 2.5 arc-minute resolution to extract (*45*). Land NPP was extracted from the SEDAC Human Appropriation of Net Primary Productivity Collection (http://sedac.ciesin.columbia.edu/es/hanpp.html) (*46*).

Ecological variables are often correlated with each other across geographic space, so we first tested the collinearity among the seven climate variables (*47*). We calculated Pearson’s correlation coefficient for each pair of variables and obtained a pairwise correlation matrix. Then, candidate variables considered for the model are discarded if they have highly collinear covariates with Pearson’s r > 0.85. This method is the most common approach in quantifying variable correlation.

Ecological variable from nearby locations tends to be more similar than one would expect from random locations (spatial autocorrelation), so various ecological variables are likely correlated with each other due to this autocorrelation. Strong spatial autocorrelation in a predictive variable can cause false results. Therefore, we first tested the spatial auto-correlation using the Global Moran Index analysis followed by Moran’s I test under randomization (*48*). We found a high spatial autocorrelation pattern across variables in all models (Moran I statistic near ±1, P values < 0.0001). For spatial models, because of the high autocorrelation between neighboring cells, we performed spatial autoregression regression models (SARMs) using the spdep package in R to explain spatial autocorrelation(*49-51*). The fitted model was presented as

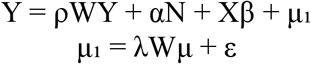

where ρ, λ and μ is the autoregression coefficient, the spatial autoregression coefficient and the spatially dependent error term, respectively; ε is the independent errors and W represents the neighborhood matrix. To reduce the computational burden based on 1° × 1° grid cells, we implemented SARMs using a Behrmann equal area WGS84 grid cells of 1.5° × 1.5°grid cells for land amniotes and 3° × 3° grid cells for marine amniotes. We identified grid cells with more than one species to conduct all subsequent analyses.

### Phylogenetic data, diversification rate estimate and extinction risk

We used a comprehensive time-calibrated species-level phylogenetic tree containing approximately 10000 bird species. A fully resolved tree overlaying on the Hackett backbone for 1,310 birds was obtained from the Bird Tree projects (*52*). For 863 mammal species, we employed a phylogenetic tree from Upham *et al*. (*53*) that consisted of a robust evolutionary timescale comprising approximately 6000 living species. We randomly extracted 1000 fully resolved trees for each clade from this supertree, and used TreeAnnotator (*54*) to build each one maximum clade credibility (MCC) tree with a burn-in of 10% of the sampled trees following previous study (*4*). For 460 reptiles, a time-calibrated phylogenetic tree was compiled from Tonini *et al*. (*55*) for squamates, from Thomson *et al*. (*56*) for turtles, and from Oaks(*57*) for crocodiles, respectively.

Diversification rate was calculated as the DR statistic (the inverse equal splits rates) which has been thought to reflect speciation rates better than net diversification rate (*52, 58*). The species-level lineage diversification rate was calculated by the following function:

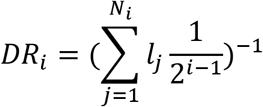

where *Ni* represents the edge number from species *j* to the root and *lj* indicates the length of the edge *j*. We determined mean diversification rates for seabird species cooccurring in each grid cell at the scale of the 1.5° × 1.5°grid cells for land amniotes and 3° × 3° grid cells for marine amniotes.

We determined extinction risks as the proportion of threatened species using two binary response variables. First, we obtained the threatened levels from the IUCN Red List (*59*) that provides the category of extinction risk for each species, including least concerned (LC), near threatened (NT), vulnerable (VU), endangered (EN), critically endangered (CR), extinct (EX), extinct in the wild (EW), and data deficient (DD). Second, extinction risk was evaluated as 1 for the categories critically endangered (CR), endangered (EN), or vulnerable (VU) and 0 for others. For spatial analyses, we determined mean extinction risks for seabird species cooccurring in each grid cell at the scale of the 1.5° × 1.5°grid cells for land amniotes and 3° × 3° grid cells for marine amniotes.

### Spatial structural equation models

We employed spatial structural equation models to examine both the direct and indirect causal effects among diversification rates, extinction risks, and variables using the R package piecewiseSEM (*60*). This method calculates path coefficients and elucidates complex multivariate relationships among a set of interrelated variables, accounting for the autocorrelation structure. We hypothesized that: 1) diversification rate is influenced by metabolic rate and environmental variables, including AMT, AMP and NPP (also including elevation for territorial patterns); 2) extinction risk is influenced by metabolic rate and environmental variables; 3) metabolic rate is influenced by body mass and environmental variables; and 4) body mass is influenced by environmental variables (see Fig. 1 B, C). The goodness of fit for the model was assessed using Fisher’s C statistics.

## Author contributions

Y.X. designed the study. Y.X., Q.Y., Z.W., and L.H. conducted the analyses and revised the manuscript. Y.X. wrote the paper.

## Data and materials availability

All data needed to evaluate the conclusions in the paper are present in the paper, in the Supplementary Materials.

