## Supplementary Information for "Global patterns, drivers and impacts of metabolic traits across amniotes"

[1] Department of Zoology, College of Life science, Sichuan Agricultural University, Ya'an 625000, China. [2] Key Laboratory of Forest Ecology in Tibet Plateau, Tibet Agricultural & Animal Husbandry University, Ministry of Education, Nyingchi 860000, China. [3] School of Life Sciences, Shandong University, Jimo District, Qingdao 266237, China. [4] Key Laboratory of Zoological Systematics and Evolution, Institute of Zoology, Chinese Academy of Sciences, Beijing 100101, China. [5] College of Life Sciences, University of Chinese Academy of Sciences, Beijing 100049, China.

[#] These authors contributed equally

**Supplementary Information includes:**

Figs. S1 to 8

Tables S1 to S4

Data S1

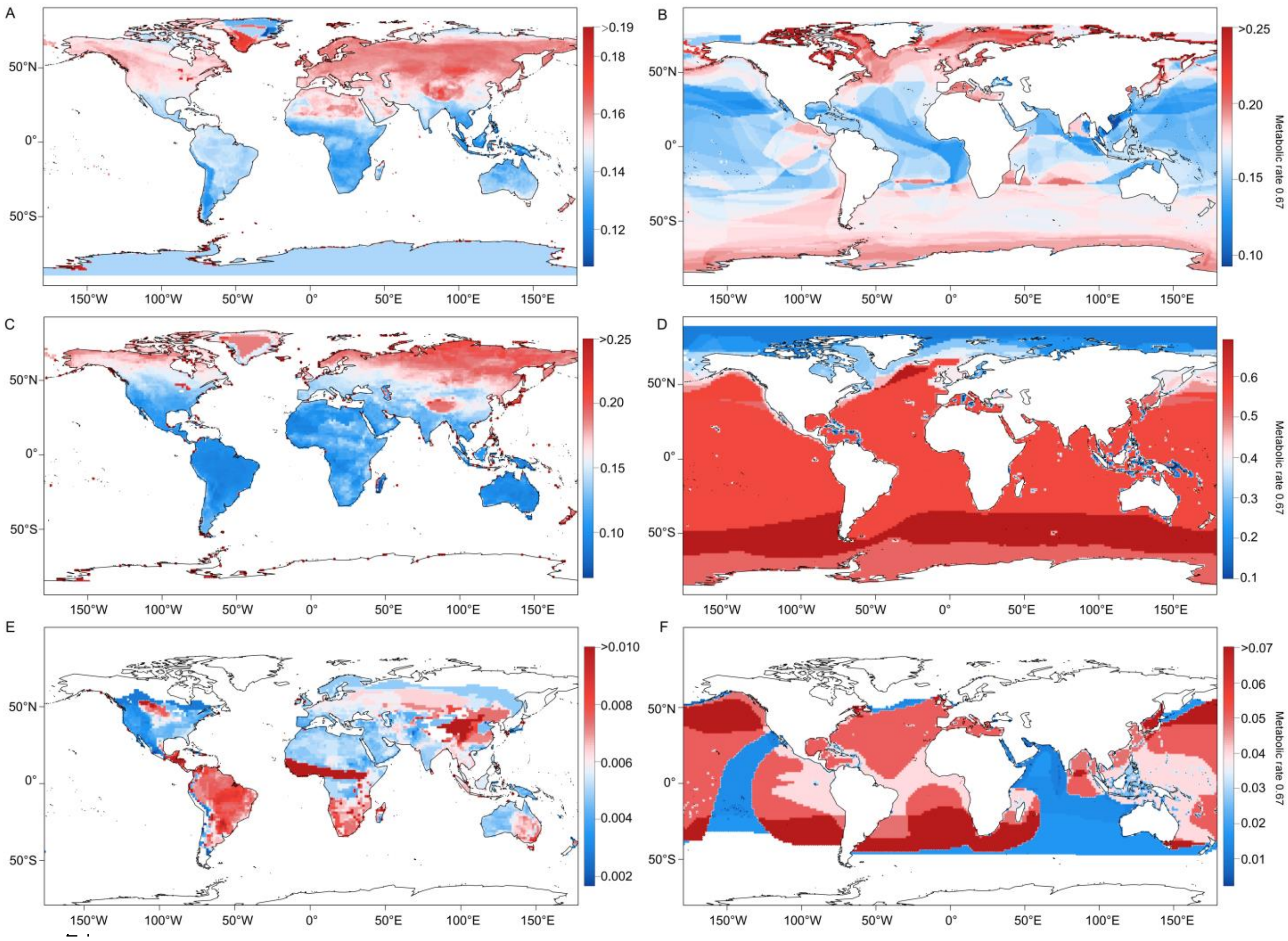

**Fig. S1 Global patterns of mass $^{2/3}$ -specific metabolic rates among terrestrial and marine amniotes.** Mass $^{2/3}$ -specific metabolic rates increase with distance from the equator across latitudes in terrestrial birds (**A**), mammals (**B**), and reptiles (**C**), while a different pattern is found among marine amniotes.

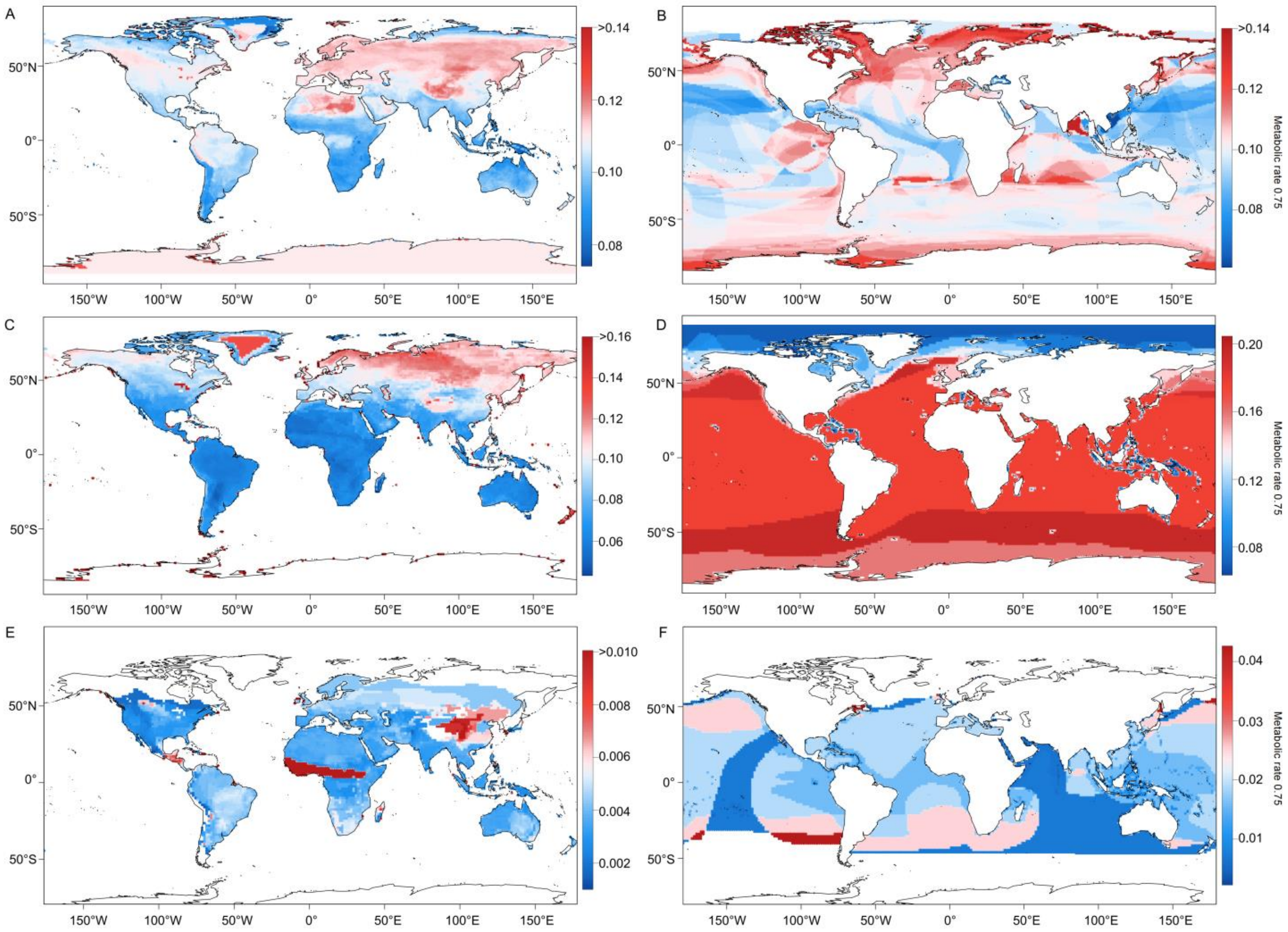

**Fig. S2 Global patterns of mass $^{3/4}$ -specific metabolic rates among terrestrial and marine amniotes.** Mass $^{3/4}$ -specific metabolic rates increase with distance from the equator across latitudes in terrestrial birds (**A**), mammals (**B**), and reptiles (**C**), while a different pattern is found among marine amniotes.

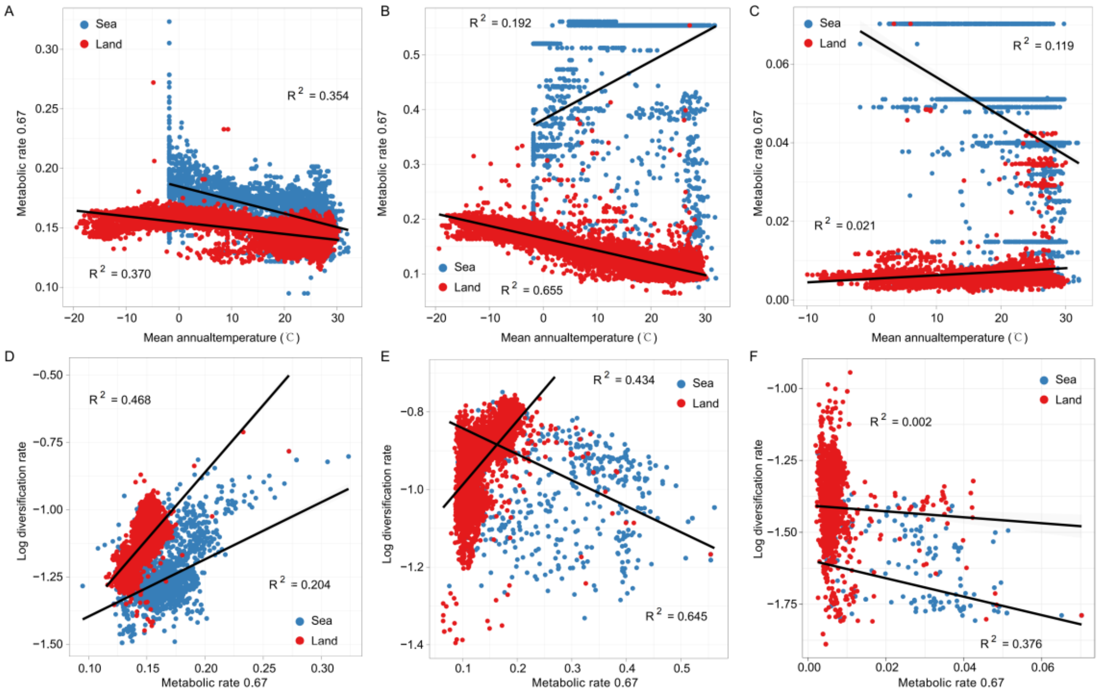

**Fig. S3 The relationships between temperature and mass $^{2/3}$ -specific metabolic rate, as well as between mass $^{2/3}$ -specific metabolic rate and diversification rate. A.** Significantly negative relationships between temperature and mass $^{2/3}$ -specific metabolic rate in birds, terrestrial mammals and marine reptiles, but positive relationships in marine mammals and land reptiles. **B.** Significantly positive relationships between relative metabolic rate and diversification rate in birds and land mammals, while negative relationships in reptiles and marine mammals.

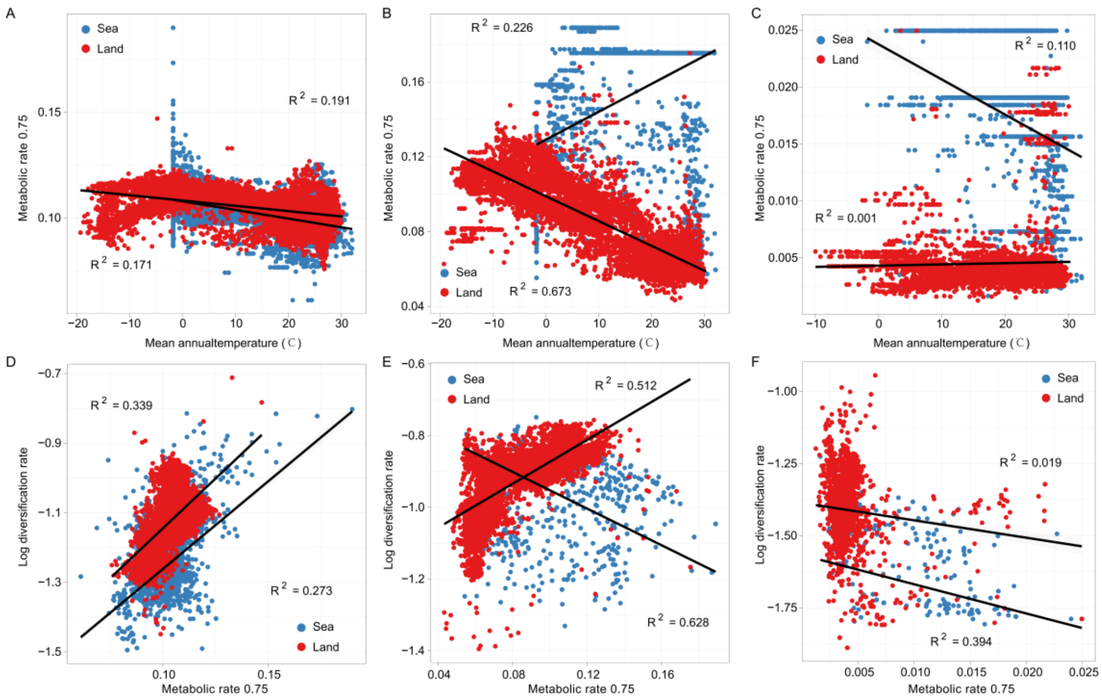

**Fig. S4 The relationships between temperature and mass $^{3/4}$ -specific metabolic rate, as well as between mass $^{3/4}$ -specific metabolic rate and diversification rate. A.** Significantly negative relationships between temperature and mass $^{3/4}$ -specific metabolic rate in birds, terrestrial mammals and marine reptiles, but positive relationships in marine mammals. **B.** Significantly positive relationships between relative metabolic rate and diversification rate in birds and land mammals, while negative relationships in reptiles and marine mammals.

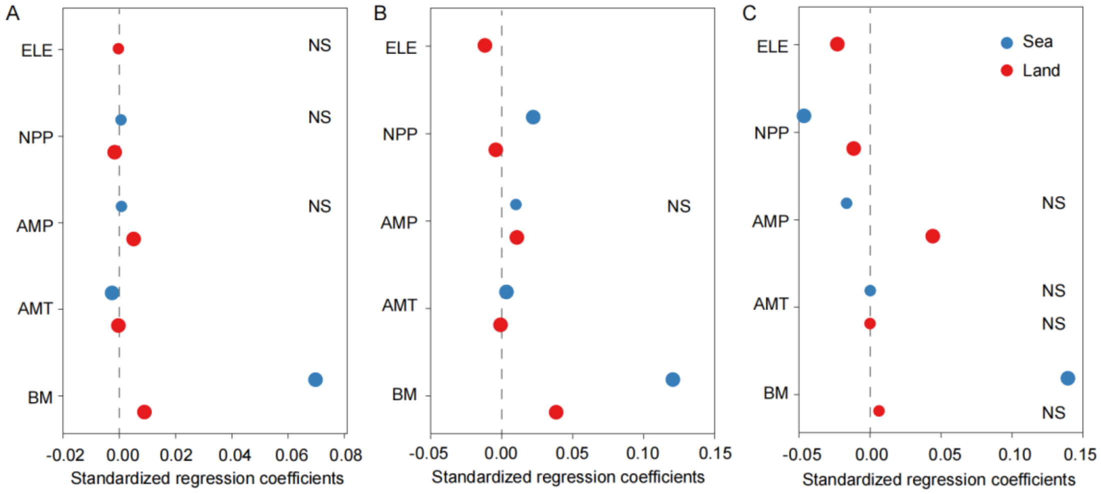

**Fig. S5 Spatial autocorrelation regression models (SARMs) on relative metabolic rates.** In birds (**A**), mammals (**B**), and reptiles (**C**). Standardized regression coefficients with standard errors in terrestrial and marine amniotes are represented by various colors. Abbreviations: AMT, annual mean temperature; AMP, annual mean precipitation; NPP, net primary productivity; ELE, elevation; BM, body mass; DR, diversification rate; ER, extinction risk.

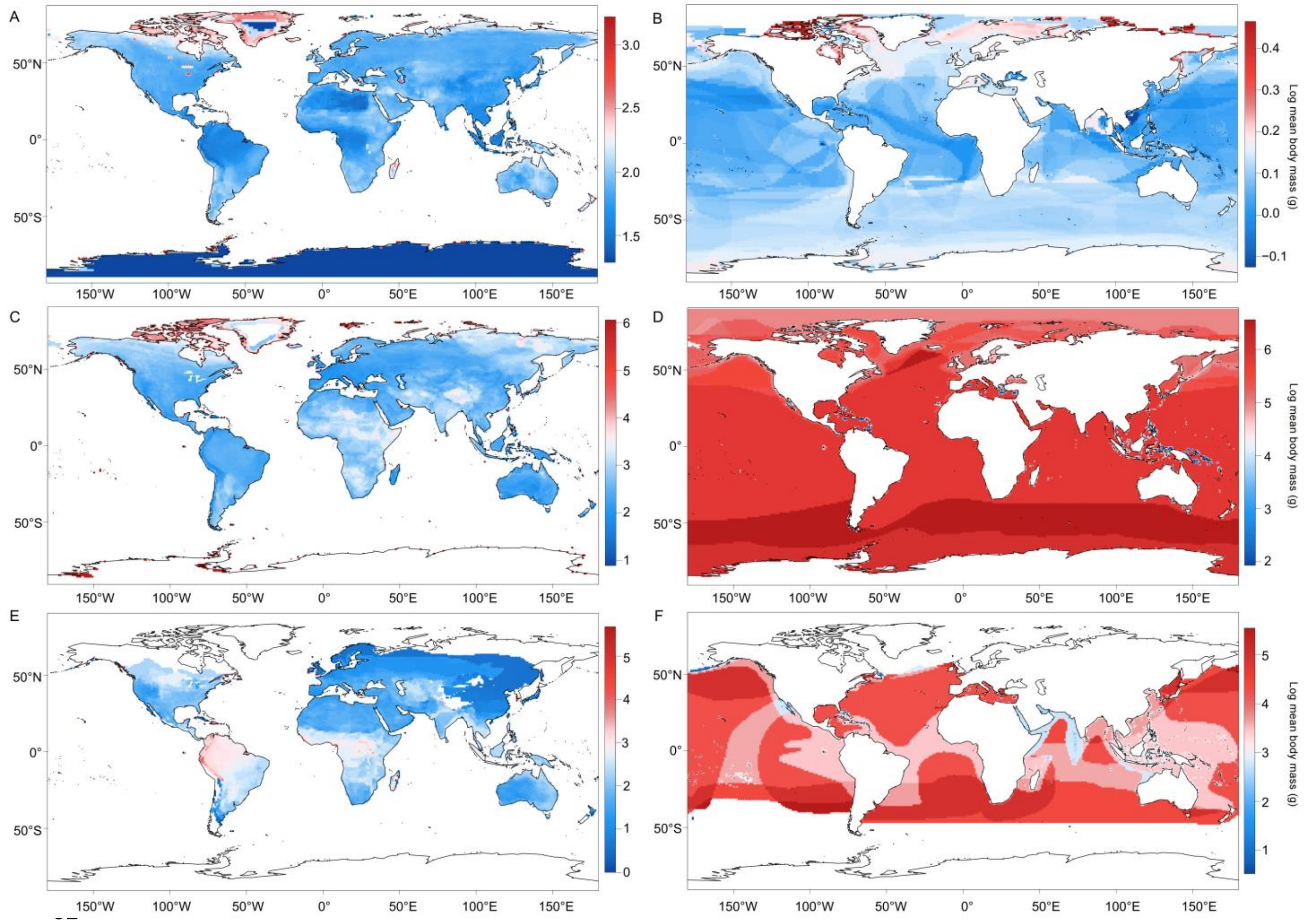

**Fig. S6 Global patterns of body mass among terrestrial and marine amniotes.** In birds (**A**), mammals (**B**), and reptiles (**C**).

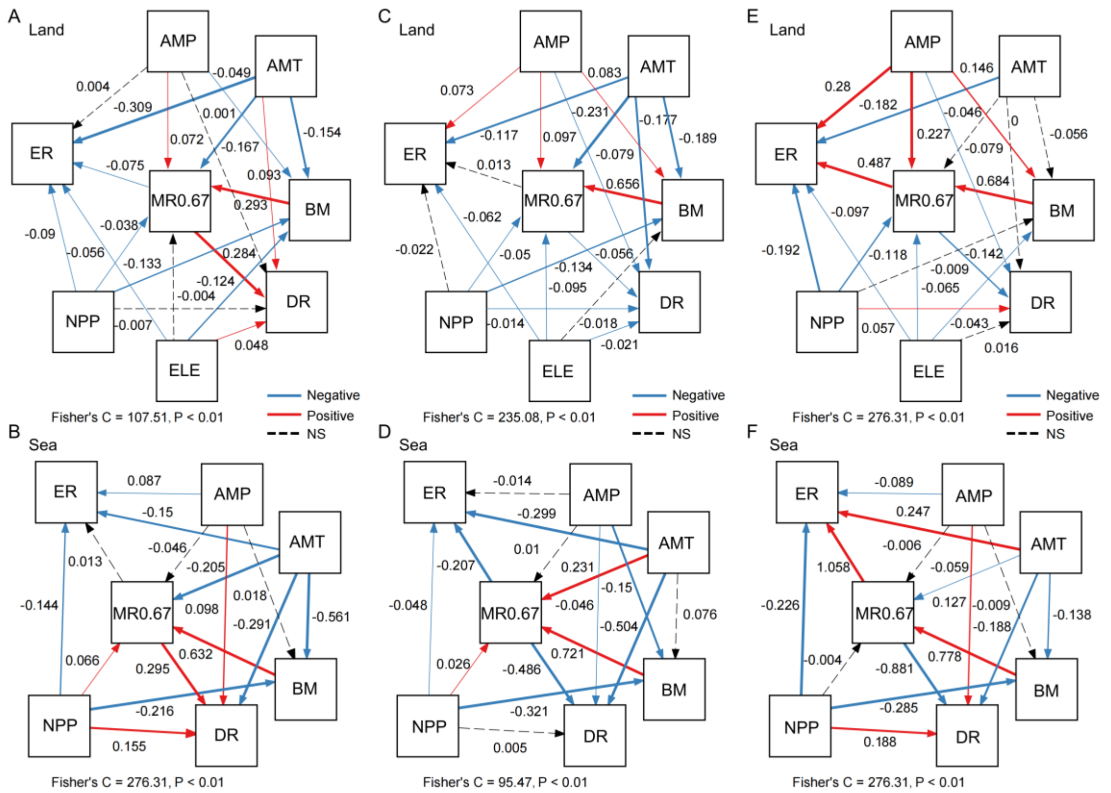

**Fig. S7 Spatial structural equation models of mass $^{2/3}$ -specific metabolic rates: drivers and impacts. A.** In birds. **B.** In mammals. **C.** In reptiles. The values on the arrows represent standardized path coefficients. Red arrows indicate significantly positive effects, while blue arrows indicate significantly negative effects. Non-significant paths are indicated by dashed arrows. The thickness of the path arrows reflects the strength of the relationships. The results of the structural equation models are summarized in Tables S3 and S4. Abbreviations: AMT, annual mean temperature; AMP, annual mean precipitation; NPP, net primary productivity; ELE, elevation; MR0.67, mass $^{2/3}$ -specific metabolic rate; BM, body mass; DR, diversification rate; ER, extinction risk.

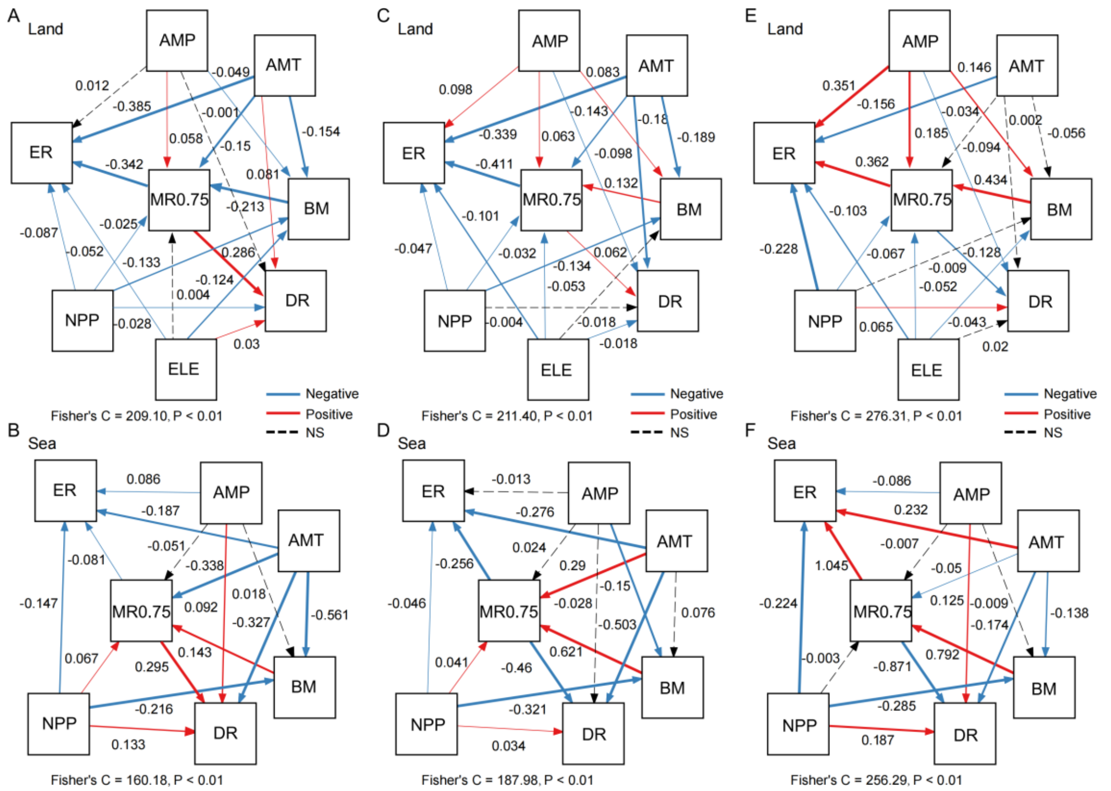

**Fig. S8 Spatial structural equation models of mass $^{3/4}$ -specific metabolic rates: drivers and impacts. A.** In birds. **B.** In mammals. **C.** In reptiles. The values on the arrows represent standardized path coefficients. Red arrows indicate significantly positive effects, while blue arrows indicate significantly negative effects. Non-significant paths are indicated by dashed arrows. The thickness of the path arrows reflects the strength of the relationships. The results of the structural equation models are summarized in Tables S3 and S4. Abbreviations: AMT, annual mean temperature; AMP, annual mean precipitation; NPP, net primary productivity; ELE, elevation; MR0.75, mass $^{3/4}$ -specific metabolic rate; BM, body mass; DR, diversification rate; ER, extinction risk.

**Table S1 Spatial autocorrelation regression models (SARMs) on metabolic rates and variables across terrestrial amniotes.**

| | Group | β | STD | z value | P |
|---|---|---|---|---|---|
| **Single-predictor SARMs** | | | | | |
| RBMR~AMT | Birds | -4.26E-05 | 6.45E-06 | -6.5997 | **4.12E-11** |
| | Mammals | -2.87E-04 | 2.71E-05 | -10.62 | **2.20E-16** |
| | Reptiles | -2.48E-04 | 6.03E-05 | -4.1043 | **4.06E-05** |
| MR0.67~AMT | Birds | -1.89E-05 | 2.52E-06 | -7.4984 | **6.46E-14** |
| | Mammals | -1.88E-04 | 1.51E-05 | -12.447 | **2.20E-16** |
| | Reptiles | 4.76E-05 | 1.97E-05 | 2.4234 | **0.0154** |
| MR0.75~AMT | Birds | -1.15E-04 | 1.25E-05 | -9.1997 | **2.20E-16** |
| | Mammals | -5.06E-05 | 5.03E-06 | -10.055 | **2.20E-16** |
| | Reptiles | 1.68E-05 | 8.68E-06 | 1.93 | 0.0536 |
| **Multivariate SARMs** | | | | | |
| **RBMR~BM + AMT + AMP + NPP + ELE** | | | | | |
| ~BM | Birds | 8.96E-03 | 1.72E-03 | 5.1974 | **2.02E-07** |
| ~AMT | Birds | -3.29E-04 | 7.03E-05 | -4.6772 | **2.91E-06** |
| ~AMP | Birds | 5.15E-03 | 9.29E-04 | 5.5401 | **3.02E-08** |
| ~NPP | Birds | -1.62E-03 | 5.16E-04 | -3.1317 | **0.001738** |
| ~ELE | Birds | -2.65E-04 | 7.71E-04 | -0.3434 | 0.731324 |
| ~BM | Mammals | 3.85E-02 | 1.47E-03 | 26.1555 | **2.20E-16** |
| ~AMT | Mammals | -7.63E-04 | 1.82E-04 | -4.199 | **2.68E-05** |
| ~AMP | Mammals | 1.08E-02 | 2.39E-03 | 4.5209 | **6.16E-06** |
| ~NPP | Mammals | -4.08E-03 | 1.32E-03 | -3.0915 | **0.001992** |
| ~ELE | Mammals | -1.18E-02 | 1.97E-03 | -5.9749 | **2.30E-09** |
| ~BM | Reptiles | 6.42E-03 | 3.72E-03 | 1.7262 | 0.0843 |
| ~AMT | Reptiles | 8.46E-05 | 5.30E-04 | 0.1596 | 0.873 |
| ~AMP | Reptiles | 4.42E-02 | 6.72E-03 | 6.5841 | **4.58E-11** |
| ~NPP | Reptiles | -1.14E-02 | 3.70E-03 | -3.0717 | **2.13E-03** |
| ~ELE | Reptiles | -2.30E-02 | 5.87E-03 | -3.9244 | **8.69E-05** |
| **MR0.67~BM + AMT + AMP + NPP + ELE** | | | | | |
| ~BM | Birds | 1.75E-02 | 6.65E-04 | 26.3002 | **2.20E-16** |
| ~AMT | Birds | -8.83E-05 | 2.67E-05 | -3.3009 | **9.64E-04** |
| ~AMP | Birds | 1.93E-03 | 3.55E-04 | 5.4445 | **5.19E-08** |
| ~NPP | Birds | -6.16E-04 | 1.98E-04 | -3.1109 | **0.0018649** |
| ~ELE | Birds | 4.78E-05 | 2.94E-04 | 0.1625 | 0.8709031 |
| ~BM | Mammals | 5.17E-02 | 7.48E-04 | 69.106 | **2.20E-16** |
| ~AMT | Mammals | -2.78E-04 | 8.51E-05 | -3.2672 | **1.09E-03** |
| ~AMP | Mammals | 7.04E-03 | 1.14E-03 | 6.1772 | **6.53E-10** |
| ~NPP | Mammals | -3.88E-03 | 6.34E-04 | -6.1205 | **9.33E-10** |
| ~ELE | Mammals | -7.36E-03 | 9.33E-04 | -7.8824 | **3.11E-15** |
| ~BM | Reptiles | 6.21E-03 | 1.95E-04 | 31.7669 | **2.20E-16** |

| | Group | β | STD | z value | P |
|---|---|---|---|---|---|
| ~AMT | Reptiles | -1.30E-05 | 2.64E-05 | -0.4913 | 0.623 |
| ~AMP | Reptiles | 2.51E-03 | 3.44E-04 | 7.2844 | **3.23E-13** |
| ~NPP | Reptiles | -1.19E-03 | 1.96E-04 | -6.0745 | **1.24E-09** |
| ~ELE | Reptiles | -1.01E-03 | 2.99E-04 | -3.375 | **7.38E-04** |
| **MR0.75~BM + AMT + AMP + NPP + ELE** | | | | | |
| ~BM | Birds | -9.46E-03 | 4.24E-04 | -22.3085 | **2.20E-16** |
| ~AMT | Birds | -5.30E-05 | 1.71E-05 | -3.0918 | **1.99E-03** |
| ~AMP | Birds | 1.16E-03 | 2.27E-04 | 5.1087 | **3.24E-07** |
| ~NPP | Birds | -3.14E-04 | 1.26E-04 | -2.4837 | **0.013** |
| ~ELE | Birds | 2.76E-04 | 1.88E-04 | 1.4683 | 0.14203 |
| ~BM | Mammals | 5.90E-03 | 3.15E-04 | 18.7054 | **2.20E-16** |
| ~AMT | Mammals | -1.14E-04 | 3.90E-05 | -2.9151 | **3.56E-03** |
| ~AMP | Mammals | 2.86E-03 | 5.14E-04 | 5.5676 | **2.58E-08** |
| ~NPP | Mammals | -1.09E-03 | 2.84E-04 | -3.8217 | **1.33E-04** |
| ~ELE | Mammals | -2.41E-03 | 4.23E-04 | -5.7043 | **1.17E-08** |
| ~BM | Reptiles | 1.89E-03 | 9.07E-05 | 20.8943 | **2.20E-16** |
| ~AMT | Reptiles | -3.86E-06 | 1.25E-05 | -0.3092 | 0.757 |
| ~AMP | Reptiles | 9.65E-04 | 1.61E-04 | 5.9869 | **2.14E-09** |
| ~NPP | Reptiles | -3.25E-04 | 9.06E-05 | -3.5841 | **3.38E-04** |
| ~ELE | Reptiles | -3.89E-04 | 1.40E-04 | -2.7688 | **5.63E-03** |

**Table S2 Spatial autocorrelation regression models (SARMs) on metabolic rates and variables across marine amniotes.**

| | Group | β | STD | z value | P |
|---|---|---|---|---|---|
| **Single-predictorSARMs** | | | | | |
| RBMR~AMT | Birds | -3.13E-04 | 3.66E-05 | -8.5426 | **2.20E-16** |
| | Mammals | 4.37E-04 | 8.87E-05 | 4.9227 | **8.54E-07** |
| | Reptiles | -0.0003 | 0.0002 | -1.8306 | 0.0672 |
| MR0.67~AMT | Birds | -1.48E-04 | 1.75E-05 | -8.4767 | **2.20E-16** |
| | Mammals | 4.23E-04 | 7.92E-05 | 5.3472 | **8.93E-08** |
| | Reptiles | -4.89E-05 | 1.85E-05 | -2.6431 | **8.22E-03** |
| MR0.75~AMT | Birds | -5.12E-04 | 6.60E-05 | -7.7625 | **8.22E-15** |
| | Mammals | 1.08E-04 | 1.97E-05 | 5.4968 | **3.87E-08** |
| | Reptiles | -1.51E-05 | 6.04E-06 | -2.5055 | **0.0122** |
| **MultivariateSARMs** | | | | | |
| **RBMR~BM+AMT+AMP+NPP+ELE** | | | | | |
| ~BM | Birds | 0.0698 | 0.0023 | 30.7073 | **0.0000** |
| ~AMT | Birds | -0.0025 | 0.0003 | -9.3762 | **0.0000** |
| ~AMP | Birds | 0.0008 | 0.0039 | 0.2093 | 0.8343 |
| ~NPP | Birds | 0.0006 | 0.0029 | 0.2011 | 0.8406 |
| ~BM | Mammals | 0.1206 | 0.0011 | 106.5684 | **0.0000** |
| ~AMT | Mammals | 0.0034 | 0.0004 | 8.1092 | **0.0000** |
| ~AMP | Mammals | 0.0100 | 0.0069 | 1.4571 | 0.1451 |
| ~NPP | Mammals | 0.0222 | 0.0051 | 4.3115 | **0.0000** |
| ~BM | Reptiles | 0.1394 | 0.0024 | 57.5681 | **0.0000** |
| ~AMT | Reptiles | 0.0002 | 0.0007 | 0.3347 | 0.7378 |
| ~AMP | Reptiles | -0.0164 | 0.0086 | -1.9056 | 0.0567 |
| ~NPP | Reptiles | -0.0465 | 0.0107 | -4.3299 | **0.0000** |
| **MR0.67~BM+AMT+AMP+NPP+ELE** | | | | | |
| ~BM | Birds | 0.0411 | 0.0011 | 39.1192 | **0.0000** |
| ~AMT | Birds | -0.0007 | 0.0001 | -6.9579 | **0.0000** |
| ~AMP | Birds | -0.0016 | 0.0017 | -0.9061 | 0.3649 |
| ~NPP | Birds | 0.0017 | 0.0013 | 1.2639 | 0.2063 |
| ~BM | Mammals | 0.1142 | 0.0007 | 160.6054 | **0.0000** |
| ~AMT | Mammals | 0.0028 | 0.0003 | 8.3818 | **0.0000** |
| ~AMP | Mammals | 0.0024 | 0.0046 | 0.5168 | 0.6053 |
| ~NPP | Mammals | 0.0134 | 0.0032 | 4.2011 | **0.0000** |
| ~BM | Reptiles | 0.0186 | 0.0002 | 89.6978 | **0.0000** |
| ~AMT | Reptiles | -0.0002 | 0.0001 | -2.0370 | **0.0417** |
| ~AMP | Reptiles | -0.0004 | 0.0008 | -0.5113 | 0.6092 |
| ~NPP | Reptiles | -0.0007 | 0.0009 | -0.7947 | 0.4268 |
| **MR0.75~BM+AMT+AMP+NPP+ELE** | | | | | |

| | Group | β | STD | z value | P |
|---|---|---|---|---|---|
| ~BM | Birds | 0.0047 | 0.0006 | 8.0548 | **0.0000** |
| ~AMT | Birds | -0.0004 | 0.0001 | -6.5532 | **0.0000** |
| ~AMP | Birds | -0.0007 | 0.0010 | -0.6849 | 0.4934 |
| ~NPP | Birds | 0.0007 | 0.0008 | 0.8546 | 0.3928 |
| ~BM | Mammals | 0.0252 | 0.0003 | 95.8683 | **0.0000** |
| ~AMT | Mammals | 0.0009 | 0.0001 | 7.5603 | **0.0000** |
| ~AMP | Mammals | 0.0010 | 0.0017 | 0.5681 | 0.5700 |
| ~NPP | Mammals | 0.0053 | 0.0012 | 4.4809 | **0.0000** |
| ~BM | Reptiles | 6.11E-03 | 7.20E-05 | 84.9597 | **0.0000** |
| ~AMT | Reptiles | -4.25E-05 | 2.61E-05 | -1.6277 | 0.104 |
| ~AMP | Reptiles | -1.80E-04 | 2.89E-04 | -0.6224 | 0.534 |
| ~NPP | Reptiles | -2.52E-04 | 3.15E-04 | -0.8025 | 0.422 |

**Table S3 Spatial structural equation models on metabolic rates and variables across terrestrial amniotes.**

| Response | Predictor | Estimate | Std. Error | Crit. Value | Std. Estimate | P | $R^2$ |
|---|---|---|---|---|---|---|---|
| **RBMR** | | | | | | | |
| **Birds** | | | | | | | |
| RBMR | BM | 0.009 | 0.002 | 5.106 | 0.046 | **0.000** | 0.93 |
| RBMR | AMT | -0.001 | 0.000 | -6.657 | -0.178 | **0.000** | |
| RBMR | AMP | 0.004 | 0.001 | 4.617 | 0.063 | **0.000** | |
| RBMR | NPP | -0.002 | 0.001 | -3.536 | -0.031 | **0.000** | |
| RBMR | ELE | -0.001 | 0.001 | -1.059 | -0.008 | 0.290 | |
| BM | AMT | -0.002 | 0.000 | -4.629 | -0.154 | **0.000** | 0.86 |
| BM | AMP | -0.017 | 0.007 | -2.666 | -0.049 | **0.008** | |
| BM | NPP | -0.042 | 0.004 | -11.035 | -0.133 | **0.000** | |
| BM | ELE | -0.064 | 0.005 | -12.323 | -0.125 | **0.000** | |
| DR | RBMR | 0.770 | 0.023 | 33.208 | 0.334 | **0.000** | 0.95 |
| DR | AMT | 0.001 | 0.000 | 4.513 | 0.100 | **0.000** | |
| DR | AMP | 0.000 | 0.002 | -0.104 | -0.001 | 0.918 | |
| DR | NPP | -0.002 | 0.001 | -2.290 | -0.016 | **0.022** | |
| DR | ELE | 0.010 | 0.002 | 6.727 | 0.042 | **0.000** | |
| ER | RBMR | -0.201 | 0.017 | -11.746 | -0.207 | **0.000** | 0.85 |
| ER | AMT | -0.001 | 0.000 | -11.085 | -0.360 | **0.000** | |
| ER | AMP | 0.000 | 0.001 | 0.348 | 0.007 | 0.728 | |
| ER | NPP | -0.006 | 0.001 | -7.454 | -0.094 | **0.000** | |
| ER | ELE | -0.006 | 0.001 | -5.997 | -0.061 | **0.000** | |
| **Mammals** | | | | | | | |
| RBMR | BM | 0.043 | 0.002 | 27.550 | 0.166 | **0.000** | 0.95 |
| RBMR | AMT | -0.001 | 0.000 | -6.731 | -0.151 | **0.000** | |
| RBMR | AMP | 0.011 | 0.003 | 4.155 | 0.047 | **0.000** | |
| RBMR | NPP | -0.005 | 0.001 | -3.258 | -0.023 | **0.001** | |
| RBMR | ELE | -0.016 | 0.002 | -7.642 | -0.048 | **0.000** | |
| BM | AMT | -0.006 | 0.001 | -5.314 | -0.189 | **0.000** | 0.79 |
| BM | AMP | 0.072 | 0.019 | 3.798 | 0.083 | **0.000** | |
| BM | NPP | -0.103 | 0.011 | -9.063 | -0.135 | **0.000** | |
| BM | ELE | -0.023 | 0.015 | -1.550 | -0.018 | 0.121 | |
| DR | RBMR | 0.104 | 0.010 | 10.509 | 0.122 | **0.000** | 0.95 |
| DR | AMT | -0.001 | 0.000 | -8.053 | -0.176 | **0.000** | |
| DR | AMP | -0.020 | 0.002 | -9.287 | -0.103 | **0.000** | |
| DR | NPP | 0.000 | 0.001 | -0.348 | -0.002 | 0.728 | |
| DR | ELE | -0.004 | 0.002 | -2.565 | -0.016 | **0.010** | |
| ER | RBMR | -0.173 | 0.015 | -11.640 | -0.323 | **0.000** | 0.72 |
| ER | AMT | -0.001 | 0.000 | -7.334 | -0.298 | **0.000** | |
| ER | AMP | 0.010 | 0.003 | 3.488 | 0.087 | **0.001** | |

| Response | Predictor | Estimate | Std. Error | Crit. Value | Std. Estimate | P | $R^2$ |
|---|---|---|---|---|---|---|---|
| ER | NPP | -0.004 | 0.002 | -2.333 | -0.041 | **0.020** | |
| ER | ELE | -0.015 | 0.002 | -6.453 | -0.089 | **0.000** | |
| **Reptiles** | | | | | | | |
| RBMR | BM | 0.002 | 0.004 | 0.473 | 0.009 | 0.636 | 0.84 |
| RBMR | AMT | -0.001 | 0.001 | -1.529 | -0.054 | 0.126 | |
| RBMR | AMP | 0.041 | 0.007 | 6.074 | 0.170 | **0.000** | |
| RBMR | NPP | -0.013 | 0.004 | -3.391 | -0.053 | **0.001** | |
| RBMR | ELE | -0.031 | 0.006 | -5.231 | -0.075 | **0.000** | |
| BM | AMT | -0.004 | 0.002 | -1.838 | -0.056 | 0.066 | 0.89 |
| BM | AMP | 0.167 | 0.027 | 6.169 | 0.146 | **0.000** | |
| BM | NPP | -0.011 | 0.015 | -0.709 | -0.009 | 0.479 | |
| BM | ELE | -0.083 | 0.024 | -3.499 | -0.043 | **0.001** | |
| DR | RBMR | -0.207 | 0.013 | -15.961 | -0.227 | **0.000** | 0.86 |
| DR | AMT | 0.000 | 0.000 | -0.264 | -0.009 | 0.792 | |
| DR | AMP | -0.019 | 0.006 | -3.405 | -0.088 | **0.001** | |
| DR | NPP | 0.014 | 0.003 | 4.295 | 0.063 | **0.000** | |
| DR | ELE | 0.004 | 0.005 | 0.779 | 0.010 | 0.436 | |
| ER | RBMR | 0.168 | 0.014 | 12.272 | 0.270 | **0.000** | 0.65 |
| ER | AMT | -0.001 | 0.000 | -2.008 | -0.085 | **0.045** | |
| ER | AMP | 0.062 | 0.006 | 11.175 | 0.413 | **0.000** | |
| ER | NPP | -0.037 | 0.004 | -10.155 | -0.238 | **0.000** | |
| ER | ELE | -0.022 | 0.005 | -4.477 | -0.086 | **0.000** | |
| **MR0.67** | | | | | | | |
| **Birds** | | | | | | | |
| MR0.67 | BM | 0.019 | 0.001 | 26.333 | 0.293 | **0.000** | 0.89 |
| MR0.67 | AMT | 0.000 | 0.000 | -5.305 | -0.167 | **0.000** | |
| MR0.67 | AMP | 0.002 | 0.000 | 4.368 | 0.073 | **0.000** | |
| MR0.67 | NPP | -0.001 | 0.000 | -3.494 | -0.038 | **0.001** | |
| MR0.67 | ELE | 0.000 | 0.000 | -0.454 | -0.004 | 0.650 | |
| BM | AMT | -0.002 | 0.000 | -4.629 | -0.154 | **0.000** | 0.86 |
| BM | AMP | -0.017 | 0.007 | -2.666 | -0.049 | **0.008** | |
| BM | NPP | -0.042 | 0.004 | -11.035 | -0.133 | **0.000** | |
| BM | ELE | -0.064 | 0.005 | -12.323 | -0.125 | **0.000** | |
| DR | MR0.67 | 2.075 | 0.055 | 37.421 | 0.284 | **0.000** | 0.95 |
| DR | AMT | 0.001 | 0.000 | 4.278 | 0.093 | **0.000** | |
| DR | AMP | 0.000 | 0.002 | 0.094 | 0.001 | 0.925 | |
| DR | NPP | -0.001 | 0.001 | -0.987 | -0.007 | 0.324 | |
| DR | ELE | 0.011 | 0.001 | 7.878 | 0.048 | **0.000** | |
| ER | MR0.67 | -0.232 | 0.043 | -5.378 | -0.075 | **0.000** | 0.85 |
| ER | AMT | -0.001 | 0.000 | -9.283 | -0.309 | **0.000** | |

| Response | Predictor | Estimate | Std. Error | Crit. Value | Std. Estimate | P | $R^2$ |
|---|---|---|---|---|---|---|---|
| ER | AMP | 0.000 | 0.001 | 0.203 | 0.004 | 0.839 | |
| ER | NPP | -0.006 | 0.001 | -7.095 | -0.090 | **0.000** | |
| ER | ELE | -0.006 | 0.001 | -5.421 | -0.056 | **0.000** | |
| **Mammals** | | | | | | | |
| MR0.67 | BM | 0.058 | 0.001 | 73.981 | 0.656 | **0.000** | 0.90 |
| MR0.67 | AMT | -0.001 | 0.000 | -8.256 | -0.231 | **0.000** | |
| MR0.67 | AMP | 0.007 | 0.001 | 6.091 | 0.097 | **0.000** | |
| MR0.67 | NPP | -0.004 | 0.001 | -5.833 | -0.062 | **0.000** | |
| MR0.67 | ELE | -0.011 | 0.001 | -11.015 | -0.095 | **0.000** | |
| BM | AMT | -0.006 | 0.001 | -5.314 | -0.189 | **0.000** | 0.79 |
| BM | AMP | 0.072 | 0.019 | 3.798 | 0.083 | **0.000** | |
| BM | NPP | -0.103 | 0.011 | -9.063 | -0.135 | **0.000** | |
| BM | ELE | -0.023 | 0.015 | -1.550 | -0.018 | 0.121 | |
| DR | MR0.67 | -0.141 | 0.015 | -9.243 | -0.056 | **0.000** | 0.95 |
| DR | AMT | -0.001 | 0.000 | -7.906 | -0.177 | **0.000** | |
| DR | AMP | -0.015 | 0.002 | -7.060 | -0.079 | **0.000** | |
| DR | NPP | -0.002 | 0.001 | -2.027 | -0.014 | **0.043** | |
| DR | ELE | -0.006 | 0.002 | -3.424 | -0.021 | **0.001** | |
| ER | MR0.67 | 0.020 | 0.025 | 0.828 | 0.013 | 0.408 | 0.71 |
| ER | AMT | -0.001 | 0.000 | -2.932 | -0.117 | **0.003** | |
| ER | AMP | 0.009 | 0.003 | 2.847 | 0.073 | **0.004** | |
| ER | NPP | -0.002 | 0.002 | -1.262 | -0.023 | 0.207 | |
| ER | ELE | -0.009 | 0.002 | -3.605 | -0.050 | **0.000** | |
| **Reptiles** | | | | | | | |
| MR0.67 | BM | 0.006 | 0.000 | 31.718 | 0.685 | **0.000** | 0.77 |
| MR0.67 | AMT | 0.000 | 0.000 | -1.140 | -0.047 | 0.254 | |
| MR0.67 | AMP | 0.002 | 0.000 | 6.910 | 0.227 | **0.000** | |
| MR0.67 | NPP | -0.001 | 0.000 | -6.275 | -0.118 | **0.000** | |
| MR0.67 | ELE | -0.001 | 0.000 | -3.852 | -0.065 | **0.000** | |
| BM | AMT | -0.004 | 0.002 | -1.838 | -0.056 | 0.066 | 0.89 |
| BM | AMP | 0.167 | 0.027 | 6.169 | 0.146 | **0.000** | |
| BM | NPP | -0.011 | 0.015 | -0.709 | -0.009 | 0.479 | |
| BM | ELE | -0.083 | 0.024 | -3.499 | -0.043 | **0.001** | |
| DR | MR0.67 | -3.011 | 0.230 | -13.110 | -0.142 | **0.000** | 0.86 |
| DR | AMT | 0.000 | 0.000 | -0.014 | -0.001 | 0.989 | |
| DR | AMP | -0.017 | 0.006 | -2.976 | -0.079 | **0.003** | |
| DR | NPP | 0.013 | 0.003 | 3.851 | 0.057 | **0.000** | |
| DR | ELE | 0.006 | 0.005 | 1.203 | 0.016 | 0.229 | |
| ER | MR0.67 | 7.063 | 0.226 | 31.214 | 0.487 | **0.000** | 0.71 |
| ER | AMT | -0.002 | 0.000 | -4.280 | -0.182 | **0.000** | |

| Response | Predictor | Estimate | Std. Error | Crit. Value | Std. Estimate | P | $R^2$ |
|---|---|---|---|---|---|---|---|
| ER | AMP | 0.042 | 0.005 | 7.794 | 0.280 | **0.000** | |
| ER | NPP | -0.030 | 0.003 | -8.925 | -0.192 | **0.000** | |
| ER | ELE | -0.024 | 0.005 | -5.270 | -0.097 | **0.000** | |
| **MR0.75** | | | | | | | |
| **Birds** | | | | | | | |
| MR0.75 | BM | -0.010 | 0.000 | -22.794 | -0.213 | **0.000** | 0.92 |
| MR0.75 | AMT | 0.000 | 0.000 | -5.497 | -0.150 | **0.000** | |
| MR0.75 | AMP | 0.001 | 0.000 | 4.144 | 0.058 | **0.000** | |
| MR0.75 | NPP | 0.000 | 0.000 | -2.807 | -0.025 | **0.005** | |
| MR0.75 | ELE | 0.000 | 0.000 | 0.468 | 0.004 | 0.640 | |
| BM | AMT | -0.002 | 0.000 | -4.629 | -0.154 | **0.000** | 0.86 |
| BM | AMP | -0.017 | 0.007 | -2.666 | -0.049 | **0.008** | |
| BM | NPP | -0.042 | 0.004 | -11.035 | -0.133 | **0.000** | |
| BM | ELE | -0.064 | 0.005 | -12.323 | -0.125 | **0.000** | |
| DR | MR0.75 | 2.811 | 0.093 | 30.137 | 0.286 | **0.000** | 0.95 |
| DR | AMT | 0.001 | 0.000 | 3.604 | 0.081 | **0.000** | |
| DR | AMP | 0.000 | 0.002 | -0.116 | -0.001 | 0.908 | |
| DR | NPP | -0.004 | 0.001 | -4.069 | -0.029 | **0.000** | |
| DR | ELE | 0.007 | 0.002 | 4.851 | 0.031 | **0.000** | |
| ER | MR0.75 | -1.418 | 0.066 | -21.518 | -0.342 | **0.000** | 0.86 |
| ER | AMT | -0.001 | 0.000 | -12.726 | -0.385 | **0.000** | |
| ER | AMP | 0.001 | 0.001 | 0.682 | 0.012 | 0.495 | |
| ER | NPP | -0.005 | 0.001 | -7.055 | -0.087 | **0.000** | |
| ER | ELE | -0.005 | 0.001 | -5.321 | -0.052 | **0.000** | |
| **Mammals** | | | | | | | |
| MR0.75 | BM | 0.007 | 0.000 | 19.986 | 0.132 | **0.000** | 0.94 |
| MR0.75 | AMT | 0.000 | 0.000 | -5.887 | -0.143 | **0.000** | |
| MR0.75 | AMP | 0.003 | 0.001 | 5.106 | 0.063 | **0.000** | |
| MR0.75 | NPP | -0.001 | 0.000 | -4.043 | -0.032 | **0.000** | |
| MR0.75 | ELE | -0.003 | 0.000 | -7.692 | -0.053 | **0.000** | |
| BM | AMT | -0.006 | 0.001 | -5.314 | -0.189 | **0.000** | 0.79 |
| BM | AMP | 0.072 | 0.019 | 3.798 | 0.083 | **0.000** | |
| BM | NPP | -0.103 | 0.011 | -9.063 | -0.135 | **0.000** | |
| BM | ELE | -0.023 | 0.015 | -1.550 | -0.018 | 0.121 | |
| DR | MR0.75 | 0.265 | 0.047 | 5.658 | 0.062 | **0.000** | 0.95 |
| DR | AMT | -0.001 | 0.000 | -8.120 | -0.180 | **0.000** | |
| DR | AMP | -0.019 | 0.002 | -8.748 | -0.098 | **0.000** | |
| DR | NPP | -0.001 | 0.001 | -0.601 | -0.004 | 0.548 | |
| DR | ELE | -0.005 | 0.002 | -2.796 | -0.018 | **0.005** | |
| ER | MR0.75 | -1.111 | 0.070 | -15.939 | -0.411 | **0.000** | 0.72 |

| Response | Predictor | Estimate | Std. Error | Crit. Value | Std. Estimate | P | $R^2$ |
|---|---|---|---|---|---|---|---|
| ER | AMT | -0.002 | 0.000 | -8.651 | -0.339 | **0.000** | |
| ER | AMP | 0.012 | 0.003 | 3.972 | 0.098 | **0.000** | |
| ER | NPP | -0.005 | 0.002 | -2.709 | -0.047 | **0.007** | |
| ER | ELE | -0.018 | 0.002 | -7.469 | -0.101 | **0.000** | |
| **Reptiles** | | | | | | | |
| MR0.75 | BM | 0.002 | 0.000 | 20.697 | 0.434 | **0.000** | 0.79 |
| MR0.75 | AMT | 0.000 | 0.000 | -0.832 | -0.034 | 0.406 | |
| MR0.75 | AMP | 0.001 | 0.000 | 5.751 | 0.185 | **0.000** | |
| MR0.75 | NPP | 0.000 | 0.000 | -3.711 | -0.067 | **0.000** | |
| MR0.75 | ELE | 0.000 | 0.000 | -3.167 | -0.052 | **0.002** | |
| BM | AMT | -0.004 | 0.002 | -1.838 | -0.056 | 0.066 | 0.89 |
| BM | AMP | 0.167 | 0.027 | 6.169 | 0.146 | **0.000** | |
| BM | NPP | -0.011 | 0.015 | -0.709 | -0.009 | 0.479 | |
| BM | ELE | -0.083 | 0.024 | -3.499 | -0.043 | **0.001** | |
| DR | MR0.75 | -5.692 | 0.528 | -10.789 | -0.128 | **0.000** | 0.86 |
| DR | AMT | 0.000 | 0.000 | 0.051 | 0.002 | 0.959 | |
| DR | AMP | -0.021 | 0.006 | -3.559 | -0.094 | **0.000** | |
| DR | NPP | 0.015 | 0.003 | 4.401 | 0.065 | **0.000** | |
| DR | ELE | 0.007 | 0.005 | 1.468 | 0.020 | 0.142 | |
| ER | MR0.75 | 10.956 | 0.541 | 20.240 | 0.362 | **0.000** | 0.67 |
| ER | AMT | -0.001 | 0.000 | -3.639 | -0.156 | **0.000** | |
| ER | AMP | 0.053 | 0.006 | 9.496 | 0.351 | **0.000** | |
| ER | NPP | -0.035 | 0.004 | -10.006 | -0.228 | **0.000** | |
| ER | ELE | -0.026 | 0.005 | -5.437 | -0.103 | **0.000** | |

**Table S4 Spatial structural equation models on metabolic rates and variables across marine amniotes.**

| Response | Predictor | Estimate | Std. Error | Crit. Value | Std. Estimate | P | $R^2$ |
|---|---|---|---|---|---|---|---|
| **RBMR** | | | | | | | |
| **Birds** | | | | | | | |
| RBMR | BM | 0.073 | 0.002 | 31.562 | 0.437 | **0.000** | 0.79 |
| RBMR | AMT | -0.002 | 0.000 | -11.000 | -0.374 | **0.000** | |
| RBMR | AMP | -0.002 | 0.003 | -0.703 | -0.016 | 0.482 | |
| RBMR | NPP | 0.007 | 0.003 | 2.405 | 0.037 | **0.016** | |
| BM | AMT | -0.016 | 0.001 | -17.439 | -0.561 | **0.000** | 0.75 |
| BM | AMP | 0.015 | 0.021 | 0.743 | 0.018 | 0.457 | |
| BM | NPP | -0.240 | 0.018 | -13.127 | -0.216 | **0.000** | |
| DR | RBMR | 0.604 | 0.026 | 23.578 | 0.321 | **0.000** | 0.79 |
| DR | AMT | -0.002 | 0.000 | -5.631 | -0.262 | **0.000** | |
| DR | AMP | 0.025 | 0.007 | 3.504 | 0.091 | **0.001** | |
| DR | NPP | 0.052 | 0.006 | 9.454 | 0.149 | **0.000** | |
| ER | RBMR | 0.073 | 0.032 | 2.333 | 0.033 | **0.020** | 0.77 |
| ER | AMT | -0.002 | 0.001 | -2.461 | -0.139 | **0.014** | |
| ER | AMP | 0.028 | 0.009 | 3.025 | 0.086 | **0.003** | |
| ER | NPP | -0.059 | 0.007 | -8.612 | -0.143 | **0.000** | |
| **Mammals** | | | | | | | |
| RBMR | BM | 0.122 | 0.001 | 111.702 | 0.699 | **0.000** | 0.92 |
| RBMR | AMT | 0.003 | 0.000 | 8.982 | 0.249 | **0.000** | |
| RBMR | AMP | 0.013 | 0.007 | 1.958 | 0.033 | 0.050 | |
| RBMR | NPP | 0.023 | 0.005 | 4.496 | 0.040 | **0.000** | |
| BM | AMT | 0.006 | 0.003 | 1.831 | 0.076 | 0.067 | 0.50 |
| BM | AMP | -0.336 | 0.074 | -4.533 | -0.150 | **0.000** | |
| BM | NPP | -1.057 | 0.070 | -15.150 | -0.321 | **0.000** | |
| DR | RBMR | -0.306 | 0.007 | -43.574 | -0.401 | **0.000** | 0.89 |
| DR | AMT | -0.006 | 0.000 | -24.316 | -0.552 | **0.000** | |
| DR | AMP | -0.010 | 0.005 | -2.082 | -0.035 | **0.037** | |
| DR | NPP | 0.014 | 0.004 | 3.175 | 0.032 | **0.002** | |
| ER | RBMR | -0.223 | 0.012 | -18.770 | -0.239 | **0.000** | 0.80 |
| ER | AMT | -0.004 | 0.001 | -6.224 | -0.296 | **0.000** | |
| ER | AMP | -0.006 | 0.010 | -0.591 | -0.017 | 0.555 | |
| ER | NPP | -0.026 | 0.008 | -3.384 | -0.049 | **0.001** | |
| **Reptiles** | | | | | | | |
| RBMR | BM | 0.140 | 0.002 | 58.140 | 0.885 | **0.000** | 0.82 |
| RBMR | AMT | 0.001 | 0.001 | 0.928 | 0.028 | 0.353 | |
| RBMR | AMP | -0.010 | 0.007 | -1.410 | -0.036 | 0.159 | |
| RBMR | NPP | -0.032 | 0.011 | -3.053 | -0.053 | **0.002** | |

| Response | Predictor | Estimate | Std. Error | Crit. Value | Std. Estimate | P | $R^2$ |
|---|---|---|---|---|---|---|---|
| BM | AMT | -0.016 | 0.006 | -2.818 | -0.138 | **0.005** | 0.68 |
| BM | AMP | -0.016 | 0.072 | -0.223 | -0.009 | 0.824 | |
| BM | NPP | -1.100 | 0.087 | -12.587 | -0.285 | **0.000** | |
| DR | RBMR | -0.701 | 0.012 | -58.411 | -0.880 | **0.000** | 0.75 |
| DR | AMT | -0.001 | 0.000 | -3.795 | -0.095 | **0.000** | |
| DR | AMP | 0.020 | 0.005 | 3.895 | 0.088 | **0.000** | |
| DR | NPP | 0.056 | 0.009 | 6.135 | 0.114 | **0.000** | |
| ER | RBMR | 1.824 | 0.022 | 81.772 | 1.040 | **0.000** | 0.83 |
| ER | AMT | 0.004 | 0.001 | 3.050 | 0.113 | **0.002** | |
| ER | AMP | -0.016 | 0.015 | -1.071 | -0.032 | 0.284 | |
| ER | NPP | -0.143 | 0.019 | -7.725 | -0.134 | **0.000** | |
| **MR0.67** | | | | | | | |
| **Birds** | | | | | | | |
| MR0.67 | BM | 0.042 | 0.001 | 40.127 | 0.632 | **0.000** | 0.73 |
| MR0.67 | AMT | 0.000 | 0.000 | -5.883 | -0.205 | **0.000** | |
| MR0.67 | AMP | -0.003 | 0.001 | -1.888 | -0.047 | 0.059 | |
| MR0.67 | NPP | 0.005 | 0.001 | 3.792 | 0.066 | **0.000** | |
| BM | AMT | -0.016 | 0.001 | -17.439 | -0.561 | **0.000** | 0.75 |
| BM | AMP | 0.015 | 0.021 | 0.743 | 0.018 | 0.457 | |
| BM | NPP | -0.240 | 0.018 | -13.127 | -0.216 | **0.000** | |
| DR | MR0.67 | 1.386 | 0.052 | 26.652 | 0.295 | **0.000** | 0.80 |
| DR | AMT | -0.003 | 0.000 | -6.388 | -0.291 | **0.000** | |
| DR | AMP | 0.027 | 0.007 | 3.819 | 0.098 | **0.000** | |
| DR | NPP | 0.054 | 0.005 | 10.014 | 0.155 | **0.000** | |
| ER | MR0.67 | 0.069 | 0.065 | 1.071 | 0.013 | 0.284 | 0.77 |
| ER | AMT | -0.002 | 0.001 | -2.659 | -0.150 | **0.008** | |
| ER | AMP | 0.028 | 0.009 | 3.030 | 0.087 | **0.002** | |
| ER | NPP | -0.060 | 0.007 | -8.676 | -0.144 | **0.000** | |
| **Mammals** | | | | | | | |
| MR0.67 | BM | 0.115 | 0.001 | 172.398 | 0.721 | **0.000** | 0.96 |
| MR0.67 | AMT | 0.003 | 0.000 | 9.082 | 0.231 | **0.000** | |
| MR0.67 | AMP | 0.004 | 0.005 | 0.774 | 0.010 | 0.439 | |
| MR0.67 | NPP | 0.014 | 0.003 | 4.296 | 0.026 | **0.000** | |
| BM | AMT | 0.006 | 0.003 | 1.831 | 0.076 | 0.067 | 0.50 |
| BM | AMP | -0.336 | 0.074 | -4.533 | -0.150 | **0.000** | |
| BM | NPP | -1.057 | 0.070 | -15.150 | -0.321 | **0.000** | |
| DR | MR0.67 | -0.406 | 0.007 | -56.461 | -0.486 | **0.000** | 0.91 |
| DR | AMT | -0.005 | 0.000 | -25.191 | -0.504 | **0.000** | |
| DR | AMP | -0.014 | 0.005 | -3.070 | -0.046 | **0.002** | |
| DR | NPP | 0.002 | 0.004 | 0.525 | 0.005 | 0.600 | |

| Response | Predictor | Estimate | Std. Error | Crit. Value | Std. Estimate | P | $R^2$ |
|---|---|---|---|---|---|---|---|
| ER | MR0.67 | -0.212 | 0.014 | -15.533 | -0.207 | **0.000** | 0.79 |
| ER | AMT | -0.004 | 0.001 | -6.269 | -0.299 | **0.000** | |
| ER | AMP | -0.005 | 0.010 | -0.485 | -0.014 | 0.628 | |
| ER | NPP | -0.026 | 0.008 | -3.310 | -0.049 | **0.001** | |
| **Reptiles** | | | | | | | |
| MR0.67 | BM | 0.019 | 0.000 | 93.540 | 0.778 | **0.000** | 0.95 |
| MR0.67 | AMT | 0.000 | 0.000 | -2.351 | -0.059 | **0.019** | |
| MR0.67 | AMP | 0.000 | 0.001 | -0.328 | -0.006 | 0.743 | |
| MR0.67 | NPP | 0.000 | 0.001 | -0.431 | -0.004 | 0.666 | |
| BM | AMT | -0.016 | 0.006 | -2.818 | -0.138 | **0.005** | 0.68 |
| BM | AMP | -0.016 | 0.072 | -0.223 | -0.009 | 0.824 | |
| BM | NPP | -1.100 | 0.087 | -12.587 | -0.285 | **0.000** | |
| DR | MR0.67 | -4.602 | 0.126 | -36.676 | -0.881 | **0.000** | 0.61 |
| DR | AMT | -0.003 | 0.001 | -4.979 | -0.188 | **0.000** | |
| DR | AMP | 0.029 | 0.008 | 3.889 | 0.127 | **0.000** | |
| DR | NPP | 0.091 | 0.012 | 7.731 | 0.188 | **0.000** | |
| ER | MR0.67 | 12.168 | 0.279 | 43.671 | 1.058 | **0.000** | 0.64 |
| ER | AMT | 0.008 | 0.002 | 4.830 | 0.247 | **0.000** | |
| ER | AMP | -0.045 | 0.021 | -2.161 | -0.089 | **0.031** | |
| ER | NPP | -0.242 | 0.027 | -9.125 | -0.226 | **0.000** | |
| **MR0.75** | | | | | | | |
| **Birds** | | | | | | | |
| MR0.75 | BM | 0.005 | 0.001 | 7.639 | 0.143 | **0.000** | 0.62 |
| MR0.75 | AMT | 0.000 | 0.000 | -8.024 | -0.338 | **0.000** | |
| MR0.75 | AMP | -0.002 | 0.001 | -1.740 | -0.051 | 0.082 | |
| MR0.75 | NPP | 0.002 | 0.001 | 3.250 | 0.067 | **0.001** | |
| BM | AMT | -0.016 | 0.001 | -17.439 | -0.561 | **0.000** | 0.75 |
| BM | AMP | 0.015 | 0.021 | 0.743 | 0.018 | 0.457 | |
| BM | NPP | -0.240 | 0.018 | -13.127 | -0.216 | **0.000** | |
| DR | MR0.75 | 2.848 | 0.105 | 27.072 | 0.295 | **0.000** | 0.80 |
| DR | AMT | -0.003 | 0.000 | -7.554 | -0.327 | **0.000** | |
| DR | AMP | 0.025 | 0.007 | 3.655 | 0.092 | **0.000** | |
| DR | NPP | 0.047 | 0.005 | 8.668 | 0.133 | **0.000** | |
| ER | MR0.75 | -0.919 | 0.131 | -7.039 | -0.081 | **0.000** | 0.77 |
| ER | AMT | -0.002 | 0.001 | -3.327 | -0.187 | **0.001** | |
| ER | AMP | 0.028 | 0.009 | 3.009 | 0.086 | **0.003** | |
| ER | NPP | -0.061 | 0.007 | -8.936 | -0.147 | **0.000** | |
| **Mammals** | | | | | | | |
| MR0.75 | BM | 0.026 | 0.000 | 102.354 | 0.621 | **0.000** | 0.93 |

| Response | Predictor | Estimate | Std. Error | Crit. Value | Std. Estimate | P | $R^2$ |
|---|---|---|---|---|---|---|---|
| MR0.75 | AMT | 0.001 | 0.000 | 9.328 | 0.290 | **0.000** | |
| MR0.75 | AMP | 0.002 | 0.002 | 1.406 | 0.025 | 0.160 | |
| MR0.75 | NPP | 0.006 | 0.001 | 4.729 | 0.041 | **0.000** | |
| BM | AMT | 0.006 | 0.003 | 1.831 | 0.076 | 0.067 | 0.50 |
| BM | AMP | -0.336 | 0.074 | -4.533 | -0.150 | **0.000** | |
| BM | NPP | -1.057 | 0.070 | -15.150 | -0.321 | **0.000** | |
| DR | MR0.75 | -1.482 | 0.031 | -47.191 | -0.460 | **0.000** | 0.90 |
| DR | AMT | -0.005 | 0.000 | -23.364 | -0.503 | **0.000** | |
| DR | AMP | -0.009 | 0.005 | -1.775 | -0.029 | 0.076 | |
| DR | NPP | 0.015 | 0.004 | 3.417 | 0.034 | **0.001** | |
| ER | MR0.75 | -1.009 | 0.055 | -18.313 | -0.256 | **0.000** | 0.80 |
| ER | AMT | -0.003 | 0.001 | -5.790 | -0.276 | **0.000** | |
| ER | AMP | -0.005 | 0.010 | -0.468 | -0.013 | 0.640 | |
| ER | NPP | -0.025 | 0.008 | -3.170 | -0.046 | **0.002** | |
| **Reptiles** | | | | | | | |
| MR0.75 | BM | 0.006 | 0.000 | 88.538 | 0.793 | **0.000** | 0.94 |
| MR0.75 | AMT | 0.000 | 0.000 | -1.973 | -0.050 | **0.049** | |
| MR0.75 | AMP | 0.000 | 0.000 | -0.379 | -0.007 | 0.705 | |
| MR0.75 | NPP | 0.000 | 0.000 | -0.319 | -0.003 | 0.750 | |
| BM | AMT | -0.016 | 0.006 | -2.818 | -0.138 | **0.005** | 0.68 |
| BM | AMP | -0.016 | 0.072 | -0.223 | -0.009 | 0.824 | |
| BM | NPP | -1.100 | 0.087 | -12.587 | -0.285 | **0.000** | |
| DR | MR0.75 | -14.070 | 0.374 | -37.613 | -0.871 | **0.000** | 0.62 |
| DR | AMT | -0.003 | 0.001 | -4.763 | -0.174 | **0.000** | |
| DR | AMP | 0.029 | 0.007 | 3.921 | 0.125 | **0.000** | |
| DR | NPP | 0.091 | 0.012 | 7.806 | 0.187 | **0.000** | |
| ER | MR0.75 | 37.156 | 0.827 | 44.943 | 1.045 | **0.000** | 0.65 |
| ER | AMT | 0.008 | 0.002 | 4.638 | 0.232 | **0.000** | |
| ER | AMP | -0.044 | 0.021 | -2.135 | -0.086 | **0.033** | |
| ER | NPP | -0.241 | 0.026 | -9.210 | -0.224 | **0.000** | |

**Data S1**

Data S1: A large dataset of metabolic rates from 2,633 species of amniote vertebrates, which includes data on 1,310 birds, 863 mammals, and 460 reptiles.
